# Behavioural and neural limits in competitive decision making: The roles of outcome, opponency and observation

**DOI:** 10.1101/571257

**Authors:** Benjamin James Dyson, Ben Albert Steward, Tea Meneghetti, Lewis Forder

## Abstract

To understand the boundaries we set for ourselves in terms of environmental responsibility during competition, we examined a neural index of outcome valence (feedback-related negativity; FRN) in relation to earlier indices of visual attention (N1), later indices of motivational significance (P3), and, eventual behaviour. In Experiment 1 (*n*=36), participants either were (*play*) or were not (*observe*) responsible for action selection. In Experiment 2 (*n*=36), opponents additionally either could (*exploitable*) or could not (*unexploitable*) be beaten. Various failures in reinforcement learning expression were revealed including large-scale approximations of random behaviour. Against unexploitable opponents, N1 determined the extent to which negative and positive outcomes were perceived as distinct categories by FRN. Against exploitable opponents, FRN determined the extent to which P3 generated neural gain for future events. Differential activation of the N1 – FRN – P3 processing chain provides a framework for understanding the behavioural dynamism observed during competitive decision making.

---

In environments where parties continually compete for mutually-exclusive outcomes, the minimization of losses and the maximization of wins is paramount (Niv, 2009). Within these contexts, loss minimization relies upon the ability to avoid exploitation. In the context of a simple, recursive non-transitive game like Rock, Paper, Scissors (RPS), the only way to guarantee the avoidance of exploitation is to operate in accordance with a mixed-equilibrium strategy (MES; Abe & Lee, 2011; Baek et al., 2013; Bi & Zhou, 2014; Loertscher, 2013). MES is characterized by selecting all three responses with equal weight across the trial series, and ensuring there are no contingencies between consecutive trials. Such behaviour renders the player unpredictable and hence unexploitable. While some species exhibit an approximation of MES behaviour under certain situations (pigeons; Sanabria & Thrailkill, 2009, and, monkeys; Lee, McGreevy & Barraclough, 2005), the demonstration of random performance in humans remains elusive (e.g., Neuringer, 1986; Terhune & Brugger, 2011). This is a result of the high cognitive demands of MES performance (Griessinger & Coricello, 2015) and the expectation that events and outcomes in the real world are not random but rather auto-correlated or ‘clumpy’ (Scheibehenne, Wilke & Todd, 2011).

One reliable precursor of when humans deviate from MES behavior- and are thus more likely to offer themselves up for exploitation-is following negative rather than positive outcomes (e.g., Dyson, Wilbiks, Sandhu, Papanicolaou & Lintag, 2016; Forder & Dyson, 2016). This predictability echoes principles of reinforcement learning such that positive outcomes are more likely to lead to behavioural repetition (*win-stay*) whereas negative outcomes are more likely to lead to behavioural change (*lose-shift*; Lee, McGreevy & Barraclough, 2005; Thorndike, 1911; Wang, Xu & Zhou, 2014). In the case of RPS, performance following negative outcomes (both *lose* and *draw* trials) lead to a decrease in *sta*y behaviour and an increase in *shift* behaviour. In contrast, performance following positive outcomes (*win* trials) tend to lead to a rough approximation of MES behaviour, but deviations from MES can be observed when the value of win is increased, making it more likely that the organism exhibits *win-stay* behavior (Forder & Dyson, 2016; see also Srihaput, Sundvall & Dyson, in preparation). Part of the reason why sub-optimal, exploitable *lose-shift* behaviour reliably occurs may be due to the self-imposed reduction in processing time allocated to decision making on trials following negative outcome (Dixon & Schreiber, 2004; Dyson, Sundvall, Forder & Douglas, 2018; Forder & Dyson, 2016; Verbruggen, Chambers, Lawrence & McLaren, 2017). Such ideas are also consistent with observations of *tilting* behaviour following loss in poker (e.g., Laakasuo et al., 2015), *chasing* behaviour following loss in roulette (e.g., Mitzenmacher & Upfal, 2005), and *post-reinforcement pauses* where gamblers revel in or ‘consume’ the current reward such that the initiation of the next trial takes longer following positive outcome (Dixon, MacLaren, Jarick, Fugelsang & Harrigan, 2013; see also Zheng et al., 2017).

To understand individual sensitivity to positive and negative outcomes at a neural level, the event-related potential (ERP) feedback-related negativity (FRN or fERN; Miltner, Brown & Coles, 1997) serves as a reliable marker. Maximal at fronto-central electrode sites and occurring approximately 200 – 300 ms after the on-set of feedback, FRN amplitudes are larger following negative relative to positive outcomes (e.g., Frank, Woroch & Curran, 2006; Gentsch, Ullsperger & Ullsperger, 2009; Holroyd, Hajack & Larsen, 2006; Nieuwenhuis, Holyord, Mol & Coles, 2004; Luft, 2014). Further modulations in FRN amplitude have also been observed between the negative and positive outcomes of others (Yeung, Holroyd & Cohen, 2005), and, as individual responsibility for outcome declines (Li, Jia, Feng, Liu, Suo & Li, 2010). Decreases in FRN difference between wins and losses for outcomes that cannot be attributed to the self (e.g., Yu & Zhou, 2006) therefore serve as a useful index regarding individual differences in the ‘motivational significance’ of feedback and the relative ownership of outcomes (Ma, Jin, Meng & Shen, 2014; Yeung et al., 2005). Consequently, FRN also served as a metric of interest in the comparisons between problem gamblers and controls (e.g., Ulrich & Hewig, in press), in an attempt to understand the characterization of problem gambling in terms of either neural hyper-(Oberg, Christie & Tata, 2011) or hypo-sensitivity (Lole, Gonsalvez & Barry, 2015).

## Experiment 1

The distinction between neural responses to culpable versus non-culpable action is critical in understanding the control of decision-making quality in competitive environments: we must know when we are responsible for our losses and try to do better but must also know when we are not responsible for our losses and not make too much of it. While reactions to negative outcome might be mitigated by believing we were not responsible for failure, we are likely to perceive control in environments outside of our sphere of influence (Langer & Roth, 1976; Clarke, 2004), and take ownership of outcomes over which we have had little or no say (see Thompson, Armstrong & Thomas, 1998, for a review). Therefore, FRN differences between positive and negative outcomes across contexts with varying degrees of responsibility provide an objective measure as to how individuals react to their current environmental state (success or failure) and the extent to which subsequent behaviour changes as a result.

In Experiment 1 we studied the ability to distinguish between outcomes for which individuals were, and were not, responsible. This was achieved by recording FRN during RPS across two counterbalanced conditions: a *play* condition where responses were selected by participants, and an *observe* condition where responses were selected for participants. After Martinez, Bonnefon & Hoskens, (2009), participants were both actively involved and had response choice in the *play* condition, and were actively involved but had no response choice in the *observation* condition, thereby reducing outcome responsibility in the latter case. We anticipated that FRN amplitudes should be maximal at fronto-central sites (e.g., Forder & Dyson, 2016; Gao, Zika, Rogers & Thierry, 2015; Li et al., 2010; Wei et al. 2015), larger for negative outcomes (*lose* and *draw*) relative to positive outcomes (*win*), and, that the difference between FRN amplitudes between losses and wins should be greater in the *play* relative to the *observe* condition (Ma et al., 2014; Yu & Zhou, 2006).

In terms of explaining individual variation in FRN difference as a function of outcome responsibility, we focused on visual attention, self-reported engagement, and, empathy. One possibility is that to reduce the motivational significance of outcomes, individuals may simply pay less attention to them. Therefore, we used the visual N1 response as an exogenous index of attention and as an early mechanism for raising neural gain on task-relevant information (Herrmann & Knight, 2001). Neural discrimination between outcomes prior to FRN have been shown in previous work at the level of a ‘frontal P2’ component (e.g., Bellebaum & Daum, 2008; Osinsky, Mussel & Hewig, 2012). However, given the nature of scalp expressed neural activity and the reversal of polarity from anterior to posterior locations, this frontal P2 likely represents an N1 at parietal sites. In terms of our predictions for Experiment 1, we expected N1 amplitude to be larger during *play* relative to *observe* conditions. However, for those individuals who gave both *play* and *observe* conditions similar attention, we expected the magnitude of FRN difference between wins and losses across the two conditions to be also similar. In other words, the difference in visual N1 between *play* and *observe* should be positively correlated with the difference between *play* and *observe* FRN. A second related possibility is that the degree to which individuals cognitively engage with outcomes increases the significance of wins and losses (Martin & Potts, 2011; Yeung et al., 2005). By assessing game engagement (Brockmeyer, Fox, Curtiss, McBroom, Burkhart & Pidruzny, 2009) at the end of both *play* and *observe* conditions, we predicted there would be a positive relationship between the difference in self-reported engagement during *play* and *observe* conditions and the neural modulation of outcome in those conditions. Finally, we considered the possibility that increased sensitivity to outcomes for which one was idea by Zhou, Yu & Zhou (2010, p. 3607) that the degree to which FRN was expressed during the observation of outcomes was “presumably through empathetic processes involved in stranger observation.” In collecting a self-report measure of empathy (Spreng, McKinnon, Mar & Levine, 2009) at the very end of the experiment, we anticipated a negative correlation between self-reported degree of empathy and FRN differences generated by *play* and *observe* conditions.

## Method

### Participants

36 individuals (29 women) from the University of Sussex participated in the study; mean age was 21.31 years (SD = 3.51) and all were right-handed. No individuals were rejected. Studies were approved for testing by the Life Sciences and Psychology Research Ethics Committee (C-REC) at the University of Sussex (ER/BJD21/4), and participants received either course credit or £20 for participation. Compensation was independent of performance within the game.

### Stimuli and apparatus

Static pictures of a white-gloved hand signaling Rock, Paper and Scissors poses were displayed center screen at approximately 6º × 6º, with participants sat approximately 57 cm away from a 22" Diamond Plus CRT monitor (Mitsubishi, Tokyo, Japan). Participants also wore a white glove. Stimulus presentation was controlled by Presentation 18.1 (build 03.31.15) and responses were recorded using a keyboard.

### Design and procedure

Participants completed 450 trials of RPS separated across 2 counterbalanced blocks (*play, observe*) of 225 trials. At the bottom of the screen, the cumulative scores for both computer (on the left) and player (on the right) were displayed, in addition to the trial count within that block. In each block, the computer played Rock, Paper and Scissors 75 times in a random order (i.e., MES). At each trial during the *play* condition, the participant pressed one of three buttons corresponding to Rock, Paper or Scissors, prompted by the presentation of a fixation cross. Due to the use of three responses in RPS, only one form of *stay* behaviour but two forms of *shift* behaviour were available: *upgrade* [select item that would have beaten your previous play] and *downgrade* [select item that would have been beaten by your previous play] (see Dyson et al., 2016, for more details).

Following their response, the participant’s selection was presented (depicted by a picture of a white glove in one of three poses) for 1000 ms. Only the participant’s selection was presented to ensure that the outcome of the trial could not be inferred by the presentation of both responses, thereby ensuring that the subsequent feedback had sufficient informational value to produce a reliable FRN (see Forder & Dyson, 2016). A black screen followed for 500 ms and feedback was then provided for a further 1000 ms center screen (i.e., *win*, *lose*, *draw*). Scores were then updated during a 500 ms period and the next trial began with a fixation cross. Participants were informed that the computer would play in a certain way and that they were to try to beat the computer across the course of the game. The *observe* condition was identical to the *play* condition, except participants initiated each trial with a fourth button and had no control over the response selection.

### Questionnaire administration

To maintain parity with a previous study in the lab (Forder & Dyson, 2016), three questionnaires were administered following the completion of each RPS block to assess individual’s degree of engagement (Brockmyer et al., 2009), the degree of anthropomorphism assigned to the computerized opponent (Epley, Akalis, Waytz & Cacioppo, Study 1; Waytz, Cacioppo & Epley, 2010, Box A1) and co-presence felt between the player and opponent (Nowak & Biocca, 2003). To assess participant empathy, the Toronto Empathy Questionnaire (TEQ; Spreng et al., 2009) was administered after all blocks of RPS had been played. Supplementary Materials A contain additional information regarding the minor changes made to these questionnaires for the purposes of the current study.

### ERP recording

Electrical brain activity was continuously digitized using a 64 channel ANT Neuro amplifier and a 1000 Hz sampling rate. Horizontal and vertical eye movements were also recorded using channels placed at the outer canthi and at inferior orbits, respectively. Data processing was conducted using BESA 5.3 Research (MEGIS; Gräfelfing, Germany). The contributions of both vertical and horizontal eye movements were reduced from the EEG record using the VEOG and HEOG artefact options in BESA. Following average referencing and using a 0.1 Hz (12 db/oct; zero phase) to 30 Hz (24 db/oct; zero phase) filter, epochs were baseline corrected according to a 200 ms pre-feedback presentation window and neural activity was examined for 800 ms post-feedback presentation. Epochs were rejected on the basis of amplitude difference exceeding 100 μV, gradient between consecutive time points exceeding 75 μV, or, signal lower than 0.01 μV, within any channel. Both N1 and FRN mean amplitude were calculated on the basis of a 50 ms window centered around the peak latency reported for each specific condition. Peak latency windows were defined as 120 – 220 ms for the N1 aggregated across ten parietal and occipital electrodes (P7, P5, PO7, PO5, O1, P8, P6, PO8, PO6, O2), and, 225 – 350 ms for the FRN aggregated across nine fronto-central electrodes (F1, Fz, F2, FC1, FCz, FC2, C1, Cz, C2; after Forder & Dyson, 2016; Ma et al, 2014).

## Results

### Behavioural data

In terms of item distribution during *play* trials, rock (35.46%) was over-played relative to paper (33.54%) and scissors (31.00%) [F(2,70) = 4.85, MSE = 0.004, *p* = .011, ƞ_p_^2^ = .121], with only the difference between rock and scissors being significant (Tukey’s HSD test, p < .05). There was no significant effect of feedback [F(2,70) = 0.74, MSE = 0.002, *p* = .482, ƞ_p_^2^ = .021], as expected from playing an opponent operating according to MES (*win* [33.94%], *lose* [32.84%], *draw* [33.22%]).

In terms of evaluating the predictability of participant strategy as a function of outcome, proportion data were entered into a two-way repeated measures ANOVA with respect to the feedback at trial *n-1* (*win*, *lose*, *draw*) and the strategy initiated at trial *n*, involving one form of repeat behaviour (*stay*) and two forms of shift behaviour (*upgrade*, *downgrade*). The main effect of strategy [F(2,70) = 7.17, MSE = 0.039, *p* = .001, ƞ_p_^2^ = .170] and interaction between feedback x strategy [F(4,140) = 7.40, MSE = 0.014, *p* < .001, ƞ_p_^2^ = .175] was significant. As shown in Figure 1, participants at a group level approximated an unexploitable, mixed-equilibrium strategy following *win* trials. However, they were less likely to *stay* and more likely to *shift* following negative outcomes. Both upgrading (39.50%) and downgrading (39.01%) were significantly different from stay (21.49%) following *lose* trials, whereas only upgrading (38.15%) was significantly different from staying (27.47%) following *draw* trials (downgrading 34.38%).

**Figure 1.**
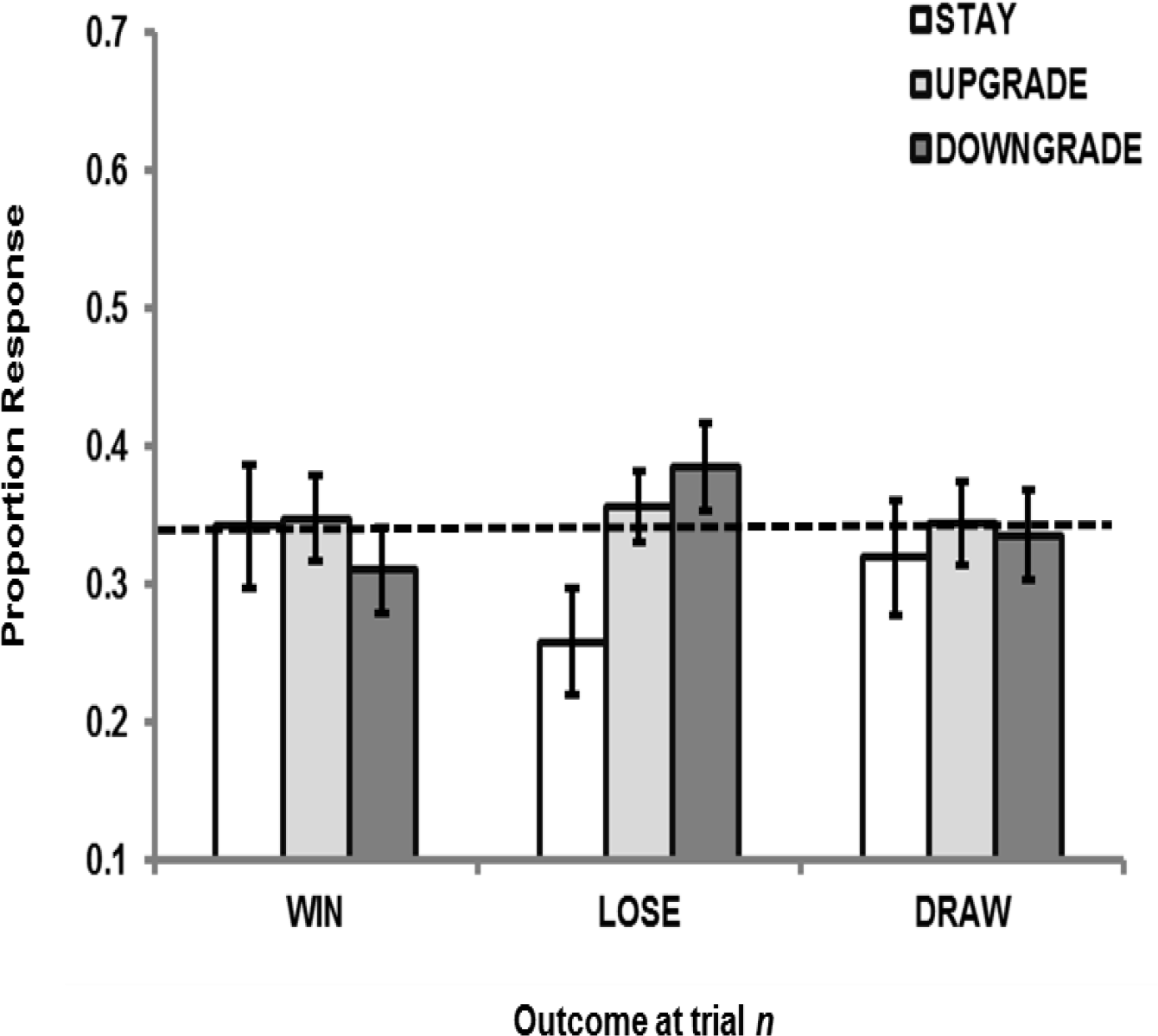
Graph showing proportion of *play n* responses in Experiment 1 separated by strategy at trial *n* (*stay, upgrade, downgrade*) as a function of outcome of trial *n-1 (win, lose, draw)* and whether trial *n-1* was a *play* or *observe* trial under a) random and b) strategic opponents. Error bars indicate +/-1 standard error and dotted line indicates a proportion of 33.3%.

### ERP data

#### Visual N1

Visual N1 was examined following the onset of feedback (*win*, *lose*, *draw*) across the two trial types (*play, observe;* see Tables 1 and 2 and Figure 1A) in terms of both peak latency and mean amplitude. A two-way repeated measures ANOVA on N1 peak latency revealed a significant main effect of feedback (*p* = .007), where the N1 peak was earlier for *draw* trials relative to *win* trials (172 vs. 177 ms; Tukey’s HSD, *p* < .05). N1 mean amplitude revealed significant main effects of trial type (*p* < .001) and feedback (*p* = .002) in the absence of a significant interaction (*p* = .122). N1 was larger during *play* than *observation* (−0.69 versus 0.63 µV), and *loss* generated smaller N1 relative to *win* and *draw* outcomes (0.21, −0.16 and −0.13 µV, respectively).

**Table 1.**
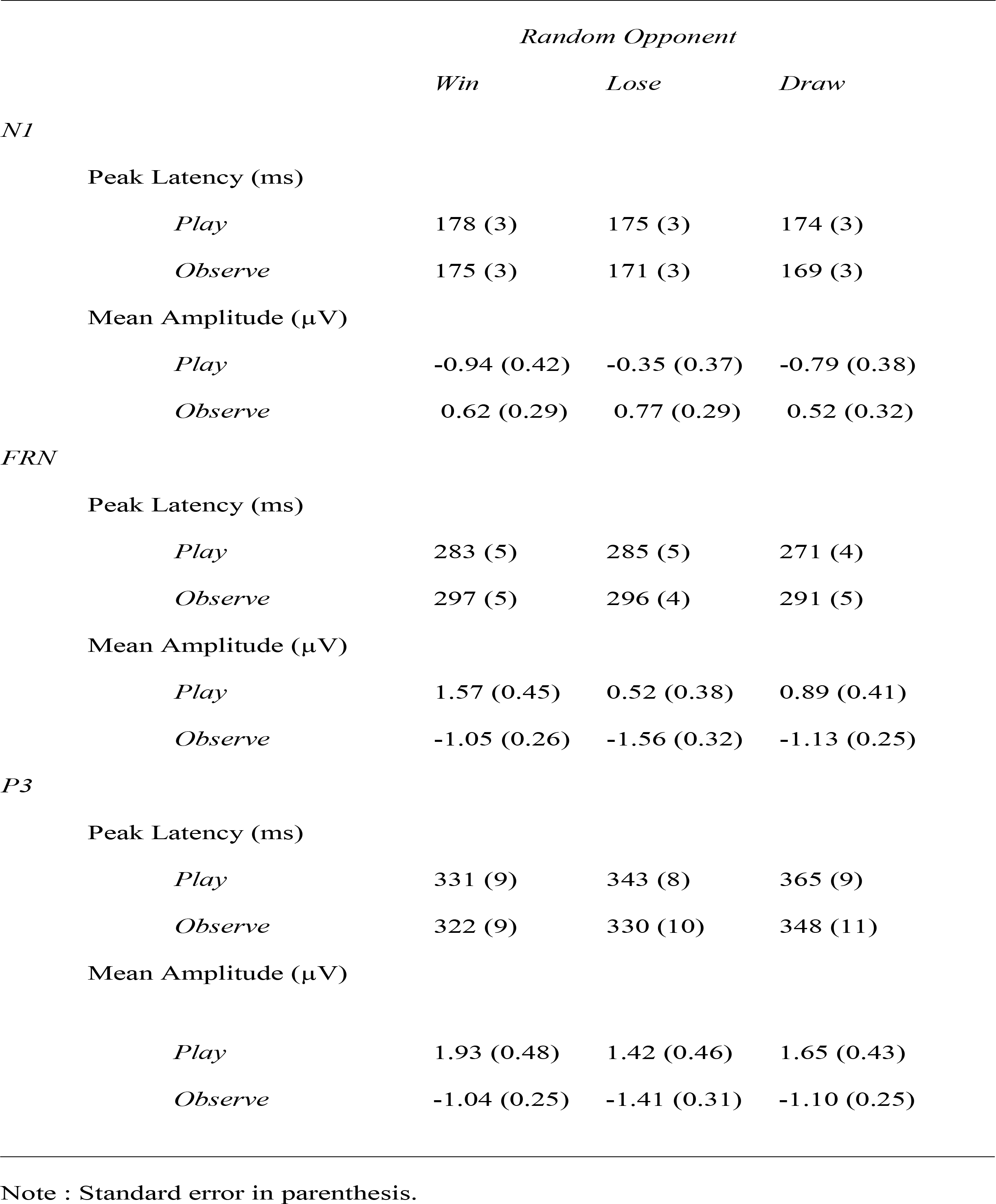
Descriptive statistics for N1, FRN and P3 components in Experiment 1

|  | *Random Opponent* |  |  |
| --- | --- | --- | --- |
|  | *Win* | *Lose* | *Draw* |
| *N1* |  |  |  |
| Peak Latency (ms) |  |  |  |
| *Play* | 178 (3) | 175 (3) | 174 (3) |
| *Observe* | 175 (3) | 171 (3) | 169 (3) |
| Mean Amplitude ($\mu\text{V}$) | | | |
| *Play* | -0.94 (0.42) | -0.35 (0.37) | -0.79 (0.38) |
| *Observe* | 0.62 (0.29) | 0.77 (0.29) | 0.52 (0.32) |
| *FRN* |  |  |  |
| Peak Latency (ms) |  |  |  |
| *Play* | 283 (5) | 285 (5) | 271 (4) |
| *Observe* | 297 (5) | 296 (4) | 291 (5) |
| Mean Amplitude ($\mu\text{V}$) | | | |
| *Play* | 1.57 (0.45) | 0.52 (0.38) | 0.89 (0.41) |
| *Observe* | -1.05 (0.26) | -1.56 (0.32) | -1.13 (0.25) |
| *P3* |  |  |  |
| Peak Latency (ms) |  |  |  |
| *Play* | 331 (9) | 343 (8) | 365 (9) |
| *Observe* | 322 (9) | 330 (10) | 348 (11) |
| Mean Amplitude ($\mu\text{V}$) | | | |
| *Play* | 1.93 (0.48) | 1.42 (0.46) | 1.65 (0.43) |
| *Observe* | -1.04 (0.25) | -1.41 (0.31) | -1.10 (0.25) |

Note : Standard error in parenthesis.

**Table 2.**
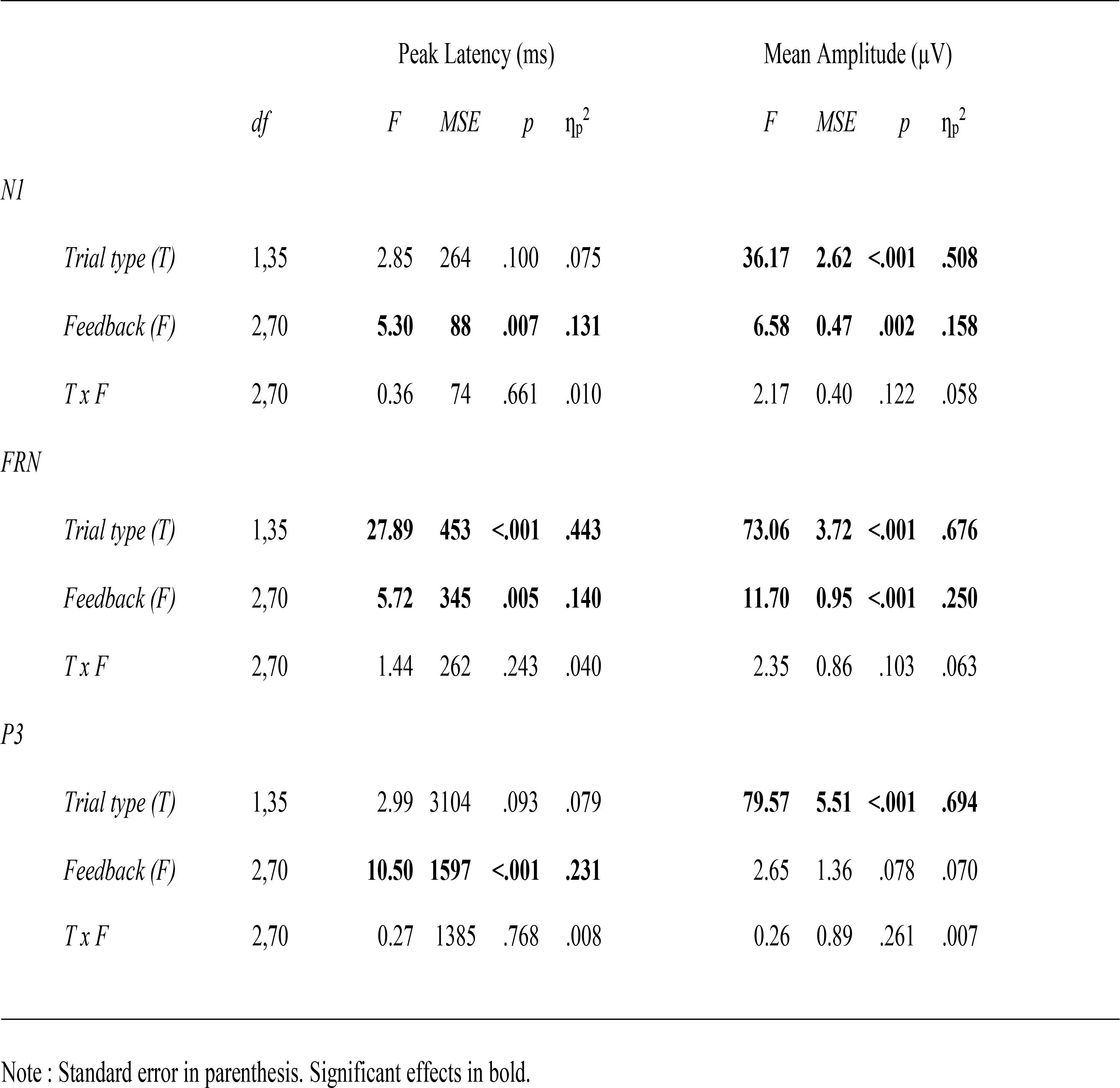
Inferential statistics for N1, FRN and P3 peak latency and mean amplitude in Experiment 1

|  |  | Peak Latency (ms) |  |  |  | Mean Amplitude (μV) |  |  |  |
| --- | --- | --- | --- | --- | --- | --- | --- | --- | --- |
|  | *df* | *F* | *MSE* | *p* | η<sub>p</sub><sup>2</sup> | *F* | *MSE* | *p* | η<sub>p</sub><sup>2</sup> |
| *N1* |  |  |  |  |  |  |  |  |  |
| *Trial type (T)* | 1,35 | 2.85 | 264 | .100 | .075 | **36.17** | **2.62** | **<.001** | **.508** |
| *Feedback (F)* | 2,70 | **5.30** | **88** | **.007** | **.131** | **6.58** | **0.47** | **.002** | **.158** |
| *T x F* | 2,70 | 0.36 | 74 | .661 | .010 | 2.17 | 0.40 | .122 | .058 |
| *FRN* |  |  |  |  |  |  |  |  |  |
| *Trial type (T)* | 1,35 | **27.89** | **453** | **<.001** | **.443** | **73.06** | **3.72** | **<.001** | **.676** |
| *Feedback (F)* | 2,70 | **5.72** | **345** | **.005** | **.140** | **11.70** | **0.95** | **<.001** | **.250** |
| *T x F* | 2,70 | 1.44 | 262 | .243 | .040 | 2.35 | 0.86 | .103 | .063 |
| *P3* |  |  |  |  |  |  |  |  |  |
| *Trial type (T)* | 1,35 | 2.99 | 3104 | .093 | .079 | **79.57** | **5.51** | **<.001** | **.694** |
| *Feedback (F)* | 2,70 | **10.50** | **1597** | **<.001** | **.231** | 2.65 | 1.36 | .078 | .070 |
| *T x F* | 2,70 | 0.27 | 1385 | .768 | .008 | 0.26 | 0.89 | .261 | .007 |

Note : Standard error in parenthesis. Significant effects in bold.

### Feedback-related negativity (FRN)

FRN was analysed in an identical manner to N1 (see Tables 1 and 2 and Figure 1B). Peak latency showed a main effect of condition (*p* < .001), indicating that FRN peak latency was earlier during *play* trials relative to *observe* trials (280 ms versus 295 ms, respectively). A main effect of feedback (*p* = .005) also showed that FRN peaked earlier during *draw* trials relative to *win* or *lose* trials (281 ms versus 290 ms and 291 ms, respectively; Tukey’s HSD test, *p* < .05). FRN mean amplitude revealed main effects of trial type (*p* < .001), feedback (*p* < .001) but no significant interaction (*p* = .103). Overall, larger positivity was generated in the *play* condition relative to the *observe* condition (0.99 µV versus −1.25 µV). With respect to feedback, we observed larger FRN amplitude for *loss* trials (−0.52 µV) relative to *win* (0.26 µV) trials (Tukey’s HSD test, *p* < .05). *Draw* trials generated intermediate FRN amplitude (−0.12 µV) and was significantly different from *win* only. Supplementary Graph A1 provides FRN difference waves between *lose* and *win* for illustrative purposes.

**Figure 2.**
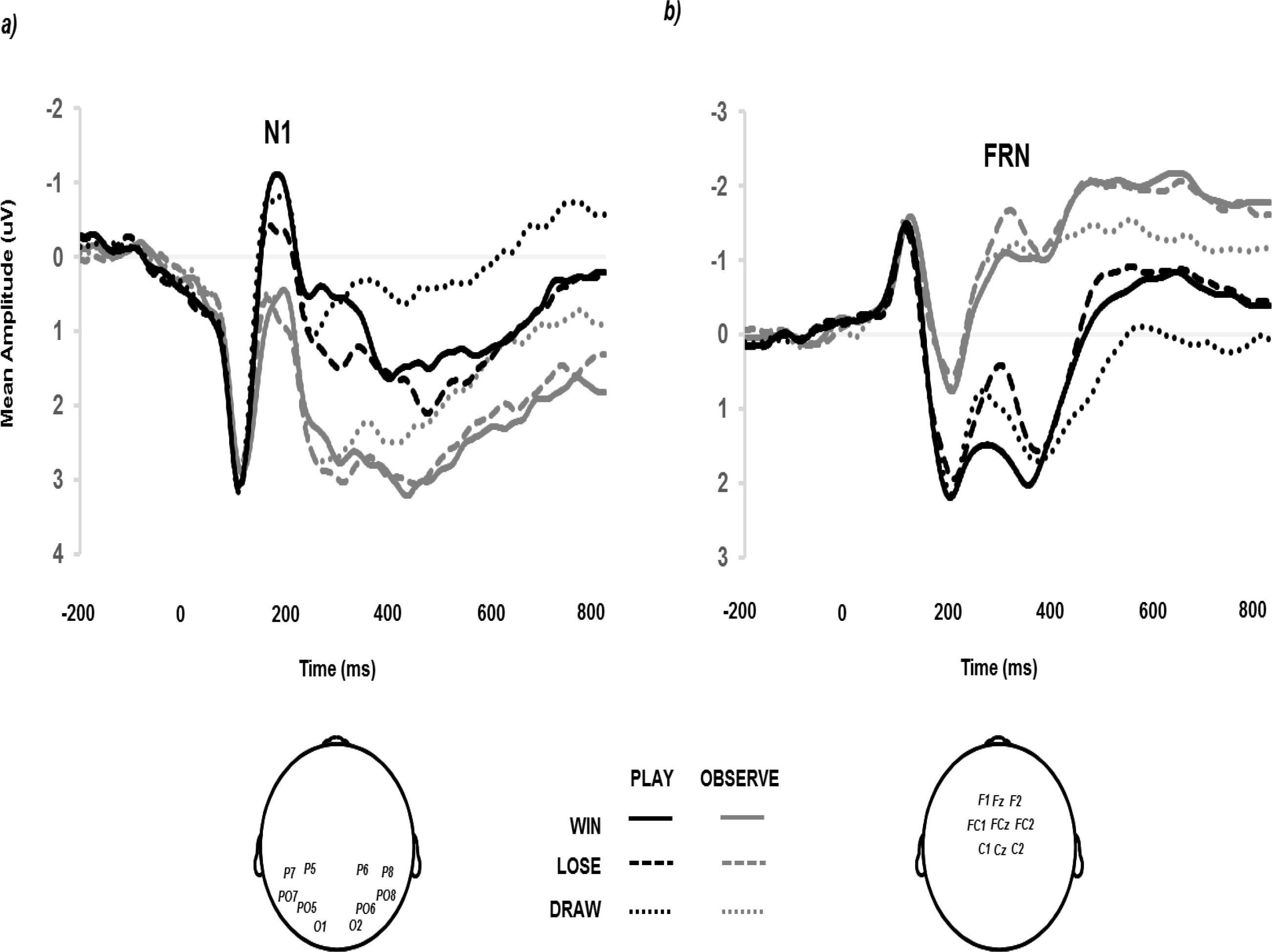
Group-average ERP from a) 10 parietal-occipital sites and b) 9 fronto-central sites generated by the presentation of trial feedback (*win, lose, draw*) and according to the nature of the trial (*play, observe*) in Experiment 1 (20 Hz filter applied for presentation).

To assess the contribution of visual attention on FRN, N1 differences between *play* and *observe* conditions [((*play win* + *play lose*) / 2) – ((*observe win* + *observe lose*) / 2)] were correlated with FRN difference between *play* and *observe* conditions [(*play lose* – *play win*) – (*observe lose* – *observe win*)] for all 36 participants. The positive correlation between N1 difference and FRN difference was significant (*r* = .373; *p* = .025; see Figure 3A). Differences in the self-reported degree of engagement between *play* and *observe* conditions failed to significantly correlate with FRN difference (*r* = -.197; *p* = .250). Finally, empathy self-report was significantly and negativity correlated with FRN difference (*r* = -.416, *p* = .012; see Figure 3B).

**Figure 3.**
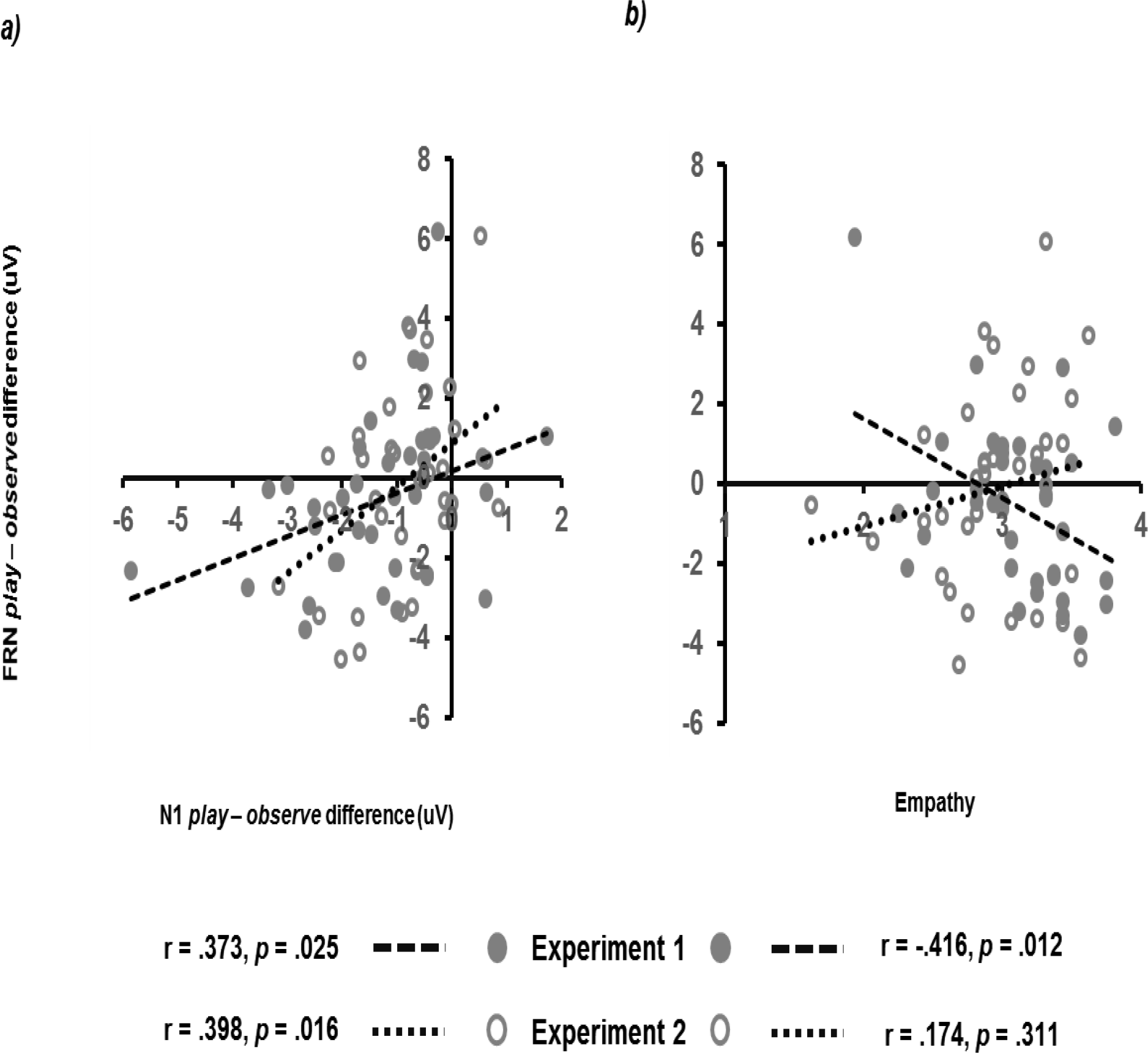
Correlations between FRN *lose – win* amplitude difference between *play – observe* conditions and a) N1 *play* – observe amplitude difference, and, *b*) self-reported empathy scores in Experiments 1 and 2.

## Discussion

At the behavioural level, RPS *play* in Experiment 1 replicated four key findings from the previous literature. First, participants showed a slight item bias towards Rock (Baek et al., 2013; Dyson et al., 2016; Forder & Dyson, 2016; Wang, Xu & Zhou, 2014). Second, participants showed an approximation of MES following positive outcomes (*win*; Dyson et al., 2016; Forder & Dyson, 2016). Third, and in contrast, participants were more predictable in their behaviour following negative (*lose*, *draw*) outcomes, specifically with respect to the expression of *shift* behaviour (Dyson et al., 2016; Forder & Dyson, 2016). Thus, performance immediately following failure is of a poorer (predictable) quality and increases the likelihood of exploitation in competitive situations.

At the neural level, we replicated the finding of larger FRN for *lose* relative to *win* outcomes at fronto-central sites (e.g., Gentsch et al., 2009). Since playing against a MES opponent yields an approximately equal distribution of outcomes, this reduces the likelihood our FRN signal was contaminated by expectancy effects (Holyroyd & Krigolson, 2007; Müller et al., 2005). Furthermore, Experiment 1 showed that this FRN difference was not statistically significant when participants both were (*play*) and were not (*observe*) responsible for response selection. This was contrary to the spirit of the previous literature where, as the motivational significance of outcomes reduce, so too does the FRN difference between those outcomes (e.g., Li et al., 2010; Ma et al., 2014; Yeung et al., 2005; Yu & Zhou, 2006). Despite this lack of difference at a group level, at an individual level there was a substantial variation in the extent to which neural difference between wins and losses was more salient when the participant was directly responsible for that outcome. The closer the degree of attention paid to the *observe* condition relative to the *play* condition (as indexed by smaller differences in visual N1 between the two conditions), the smaller the difference in *loss – win* FRN between *observe* and *play* conditions. Therefore, whereas self-report measures of attention may not predict outcome sensitivity (Yeung et al., 2005), objective neural measures such as visual N1 may help our understanding of the contribution of early attention on evaluating the importance of outcomes further downstream. The N1 data were also interesting in terms of their connections with previous findings showing that the location of a losing move in both RPS and arm wrestling appears to capture attention (Sun, Bai, Yu, Zhou, Zhang & Shen, 2015). The main effect of *smaller* visual N1 for losses relative to wins would suggest the opposite-that at a very early stage participants are ready to move on from the failure of the previous trial (e.g., Dixon et al., 2013). There was also some evidence to suggest that empathy blurred the neural distinction between outcomes for which one was (*play*) and was not (*observe*) directly responsible (Fukushima & Hiraki, 2006). That is, the higher the self-reported level of empathy (TEQ; Spreng et al., 2008), the smaller the difference between the FRN generated for losses and win across *play* and *observe* conditions.

One reason for failing to find clear group differences between *play* and *observe* trials in Experiment 1 was because the manipulation of responsibility took place at a tonic (i.e., block-by-block) level (Martin & Potts, 2011). Therefore, in the interests of replicating and extending our findings, responsibility was manipulated at a phasic (i.e., trial-by-trial) level, and, studied both in the context of an unexploitable (as per Experiment 1) and exploitable opponent in Experiment 2.

## Experiment 2

One reason for assuming that the effects associated with outcome responsibility might be better highlighted at a phasic rather than tonic level is due to the nature of neurotransmitter production associated with the motivational significance of *play* versus *observe* trials. While FRN variation is largely associated with top-down suppression of dopamine release in the basal ganglia projecting to ACC (Alexander, DeLong & Strick, 1986), norepinephrine, serotonin, GABA, and adenosine are also thought to be involved (Frank et al., 2006; Holroyd et al., 2006; Nieuwenhuis et al., 2004; Luft, 2014). Dopamine modulation is also implicated in the expression of the further down-stream frontal P3a component (Polich, 2007). ACC activity also impacts on the locus coeruleus (LC) where the release of phasic norepinephrine is putatively a causal factor in the generation of the parietal P3b (Nieuwenhuis, Aston-Jones & Cohen, 2005). Specifically, it is proposed that the ACC generates requests for increased resources on the basis of the current stimulus, and these requests are fed to the LC that subsequently increases arousal and raise the gain on task-relevant information for future stimuli (Olivera, McDonald & Goodman, 2007). This strongly suggests the hypothesis that the degree of sensitivity exhibited by the FRN in terms of differential *win* and *lose* responses may be similarly reflected in the P3 due to dopaminergic and / or norepinephrine influence.

With Experiment 2 providing the opportunity for response choice (*play*) and non-response choice (*observe*) encounters with both unexploitable (*random*) and exploitable (*strategic*) opponents, we can make the following predictions regarding motivational significance and P3. In terms of the nature of the trial, because response choice is removed during *observe* trials, this should generate smaller P3 than *play* trials. In terms of the nature of opponency, when there is nothing that can be done to improve performance as when playing against an unexploitable (*random*) opponent, each event on average should be less motivationally salient relative to the potential development of a mental model of exploitation against a *strategic* opponent. Hence, P3 should also be smaller during *random* relative to *strategic* conditions.

However, given that P3 is sensitive to both motivational significance and stimulus frequency (Nieuwenhuis et al., 2005; Polich, 2003), it will be important to directly address the contribution of the second factor in the current empirical context. For example, a recent study by Fielding et al. (2018) showed that the P3 was differentially sensitive to wins and losses. While the authors claimed that this could be taken as evidence for the P3 as representing the “brain’s reward response” they were also clear in stating that when P3 differed between outcomes, there was also a difference in the probability of winning and losing within the block (see Dyson, Forder & Sundvall, 2018, for more detailed discussion). Therefore, since P3 amplitudes are in part driven by the relevant frequency of the events to which they refer (e.g., Zheng et al, 2017), then we should additionally observe a positive correlation between the difference in *win* – *lose* rate during *strategic* conditions and the difference in P3 amplitude between *lose* and *win* trials.

Finally, to return to the issue of optimal behaviour during competition, we consider what impact *observe* trials may have on *play* trials when they are randomized within the same block. Against an unexploitable opponent and in the context of continuous *play*, we have previously observed that there is a preponderance of *shift* behavior following failure and this increase in predictability (and hence potential exploitability) may be due to individuals being more impulsive following negative outcome (Dyson et al., 2016; Dyson et al., 2018; Forder & Dyson, 2016; Verbruggen et al., 2017). If *observe* trials serve to break the cycle of poor quality *play* behaviour against *unexploitable* opponents, then *observe-play* trial pairs might help to reduce the degree to which individuals place themselves in exploitable positions by enabling a regression to MES, relative to *play-play* trial pairs. However, it is clear that in other contexts this hypothesized regression will be disadvantageous. As previously mentioned, success in competition not only relies on the minimization of losses via the ability to avoid exploitation, but also requires the maximization of wins via the ability to exploit others. In terms of successfully learning counter-strategies within RPS (e.g., Stöttinger, Filipowicz, Danckert & Anderson, 2014), *observe* trials might compromise performance by introducing an extended period across which irrelevant information (in terms of response and outcome) needs to be suppressed, while relevant information related to strategy and counter-strategy during *play* trials needs to be retained. Therefore, *observe-play* trial pairs relative to *play-play* trial pairs might also reduce the degree to which individuals express the correct strategy against *exploitable* opponents.

## Method

36 individuals (20 women, 4 left-handed) took part in the study, with a mean age of 23.94 years (SD = 6.35). Ethical approval was granted by the Life Sciences and Psychology Research Ethics Committee (C-REC) at the University of Sussex under the protocol (ER/BS300/1). All participants gave their informed consent for the study, and all were entered into a prize draw to win £25 for their participation.

Stimuli and apparatus were as per Experiment 1. Participants completed a total of 480 trials of RPS split into two, counterbalanced opponent conditions (*random, strategic*). Each opponent condition was further divided into two blocks of 120 trials. In each *random* block, the computerized opponent played according to MES, selecting 40 Rock, Paper and Scissors responses in random order. In each *strategy* block, the opponent played 80% of the time (96 trials per block) in accordance to a rule that required as the winning response a form of *shift* behavior where participants *downgraded* their previous selection, irrespective of the outcome of the previous trial. For the remaining 20% of the time (24 trials per block), MES was followed. Each block was further divided into 60 *play* and 60 *observe* trials, in a randomized order. Participants were informed at the start of each block if the computer opponent was going to play randomly or strategically.

The basic procedure of Experiment 2 was identical to Experiment 1 (see Supplementary Materials B for additional details and results). All ERP parameters were also the same as Experiment 1. The only addition was the measurement of P3a taken at the same 9 electrode sites as FRN across a 50 ms time window defined by individual condition peak latencies between 250 - 500 ms post-outcome on-set.

## Results

### Behavioural Data

#### Item selection and outcome

Proportion data during *play* trials was first analysed according to opponent (*random*, *strategic*) as a function of item selection (*rock*, *paper*, s*cissors*), and, feedback (*win*, *lose*, *draw*) in two separate, repeated-measures ANOVAs. In terms of item selection, there was no main effect of item [F(2,70) = 0.99, MSE = 0.185, *p* = .376, ƞ_p_^2^ = .028], nor an interaction with condition [F(2,70) = 1.06, MSE = 0.006, *p* = .351, ƞ_p_^2^ = .029]. There was a numerical tendency to overplay Rock (34.88%) relative to Paper (31.69%) or Scissors (33.42%), consistent with Experiment 1. In terms of feedback, there was a significant main effect [F(2,70) = 31.34, MSE = 0.011, *p* < .001, ƞ_p_^2^ =.472] and interaction with condition [F(2,70) = 38.50, MSE = 0.011, *p* < .001, ƞ_p_^2^ =.524]. This interaction confirmed the equivalence of *win*, *lose* and *draw* outcomes during the *random* condition (32.66%, 33.36%, 33.98%, respectively) as would be expected on the basis of playing an MES opponent, as per Experiment 1. Performance during the *strategic* condition confirmed that learning had taken place at a group level due to a larger proportion of wins relative to losses and draws (49.63%, 21.62%, 28.75%, respectively).

#### Random opponent performance

Proportion data from *play* trials against a *random* opponent were analysed according to trial *n-1* type (*play, observe*), feedback at trial *n-1* (*win, lose, draw*) and strategy at trial *n* (*stay, upgrade, downgrade*) in a three-way repeated-measures ANOVA (see left panel of Figure 4). There was no main effect of strategy [F(2,70) = 0.46, MSE = .103, *p* = .633, ƞ_p_^2^ = .013], no interaction between strategy x Behavioural and neural limits in competitive decision making trial *n-1* type [F(2,70) = 0.25, MSE = .073, *p* = .777, ƞ_p_^2^ = .007], no main effect of feedback at trial *n-1* [F(4,140) = 1.78, MSE = .026, *p* = .136, ƞ_p_^2^ = .048], nor was there a three-way interaction [F(4,140) = 0.99, MSE = .021, *p* = .416, ƞ_p_^2^ = .027]. As is clear from Figure 4, participants performed approximately in accordance with MES during *play* trials, and did so to an equivalent degree irrespective of whether the preceding trial *n-1* was a *play* or *observe* trial. Therefore, and contrary to Experiment 1, there was no evidence of *lose-shift* behaviour in Experiment 2. Furthermore, there was no evidence that participants were more likely to regress to MES *play* behaviour following an *observe* trial relative to a *play* trial.

**Figure 4.**
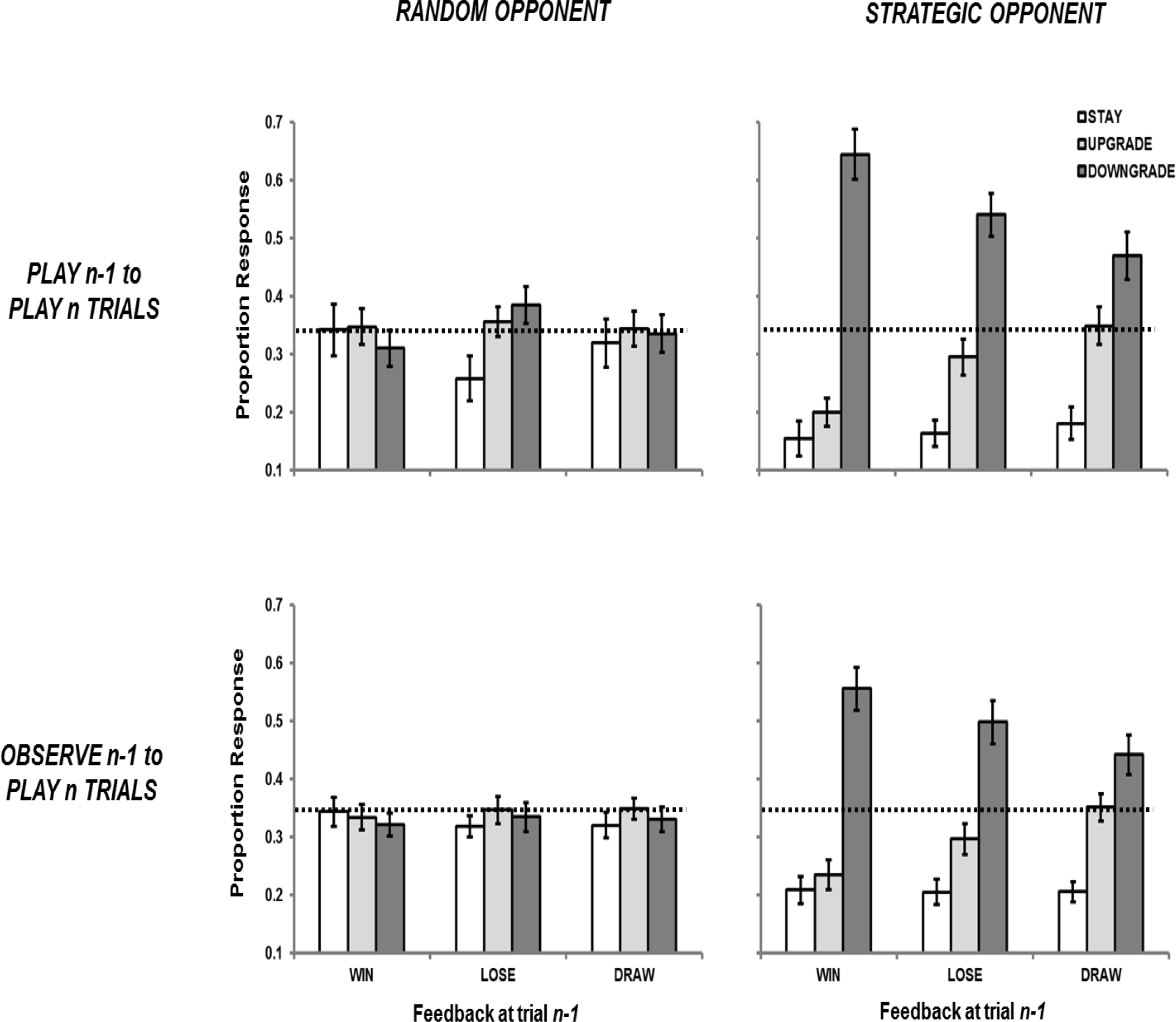
Graph showing proportion of *play n* responses in Experiment 2 separated by strategy at trial *n* (*stay, upgrade, downgrade*) as a function of outcome of trial *n-1 (win, lose, draw)* and whether trial *n-1* was a *play* or *observe* trial against *random* and *strategic* opponents. Error bars indicate +/-1 standard error and dotted line indicates a proportion of 33.3%.

#### Strategic opponent performance

Proportion data from *play* trials against a *strategic* opponent were analysed in an identical way to the previous analysis (see right panel of Figure 4). A main effect of strategy [F(2,70) = 37.23, MSE = .176, *p* < .001, ƞ_p_^2^ = .515] was subsumed in a two-way interaction with trial *n-1* feedback [F(4,140) = 14.37, MSE = 0.024, *p* < .001, ƞ_p_^2^ = .291], and a two-way interaction with trial *n-1* type [F(2,70) = 3.89, MSE = 0.032, *p* = .025, ƞ_p_^2^ = .100]. The three-way interaction between trial *n-1* type x feedback at trial *n-1* x strategy at trial *n* was not significant [F(4,140) = 0.70, MSE = 0.021, *p* = .595, ƞ_p_^2^ = .020].

In terms of the interaction between trial *n-1* feedback and trial *n* strategy, the utilization of the correct strategy (*downgrade*) fell when the previous trial was a *lose* or *draw* trial relative to a *win* trial (*win* = 60.03%, *lose* = 51.96%, *draw* = 45.64%; all differences Tukey’s HSD test, *p* < .05). In contrast, the utilization of the incorrect strategy *upgrade* rose when the previous trial was a *lose* or *draw* trial relative to a *win* trial (*win* = 21.78%, *lose* = 29.61%, *draw* = 35.03%), whereas the *stay* strategy remained relatively constant (*win* = 18.19%, *lose* = 18.44%, *draw* = 19.33%). Therefore, the outcome of the previous trial impacted on the ability to make the optimal selection for the current trial, with both forms of negative feedback (*lose* and *draw*) reducing the *downgrade* option.

Regarding the interaction between trial *n-1* type and trial *n* strategy, and similar to the previous interaction, the optimal *downgrade* option fell when the previous trial was an *observe* relative to a *play* trial (49.90% vs. 55.19%; Tukey’s HSD test, *p* < .05), whereas *staying* and *upgrading* rose when the previous encounter was an *observe* trial (16.64% vs. 20.66%, and, 28.17% vs. 29.44%, respectively), although no comparisons were significant (all Tukey’s HSD test, *p* > .05). Therefore, a previous *observe* trial reduced optimal responding on the current trial, relative to a previous *play* trial.

### ERP data

#### Visual N1

Peak latency analysis was conducted using a three-way repeated-measures ANOVA including opponent (*random, strategic*), trial *n* type (*play, observe*) and trial *n* feedback (*win, lose, draw*; see Tables 3 and 4, and, Figure 5). Main effects of opponent (*p* = .034) and trial type (*p* = .047) showed faster N1 peak latency during *strategic* relative to *random* opponents (178 versus 180 ms), and, during *observe* relative to *play* trials (178 versus 180 ms). Similarly conducted mean amplitude analysis revealed main effects of opponent (*p* = .034), trial type (*p* < .001) and feedback (*p* < .001). N1 was larger for *strategic* relative to *random* opponents (−0.21 versus 0.14 µV), larger for *play* relative to *observe* trials (−0.55 versus 0.47 µV), and, larger for *wins* relative to both *losses* and *draws* (−0.35, 0.16 and 0.08 µV, respectively; Tukey’s HSD, *p* < .05). The effects of trial type and feedback in random and strategic opponents replicate those shown in Experiment 1.

**Table 3.**
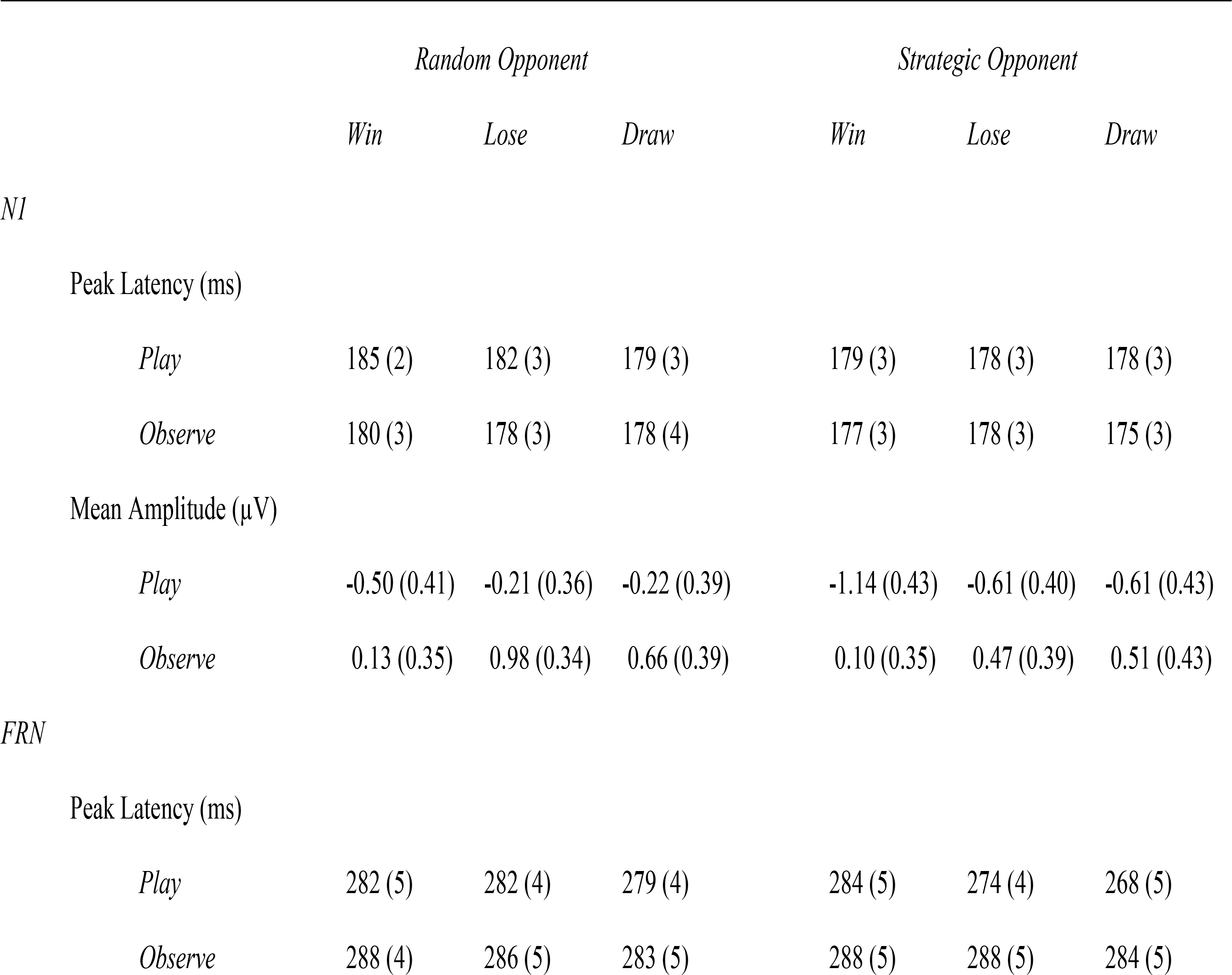
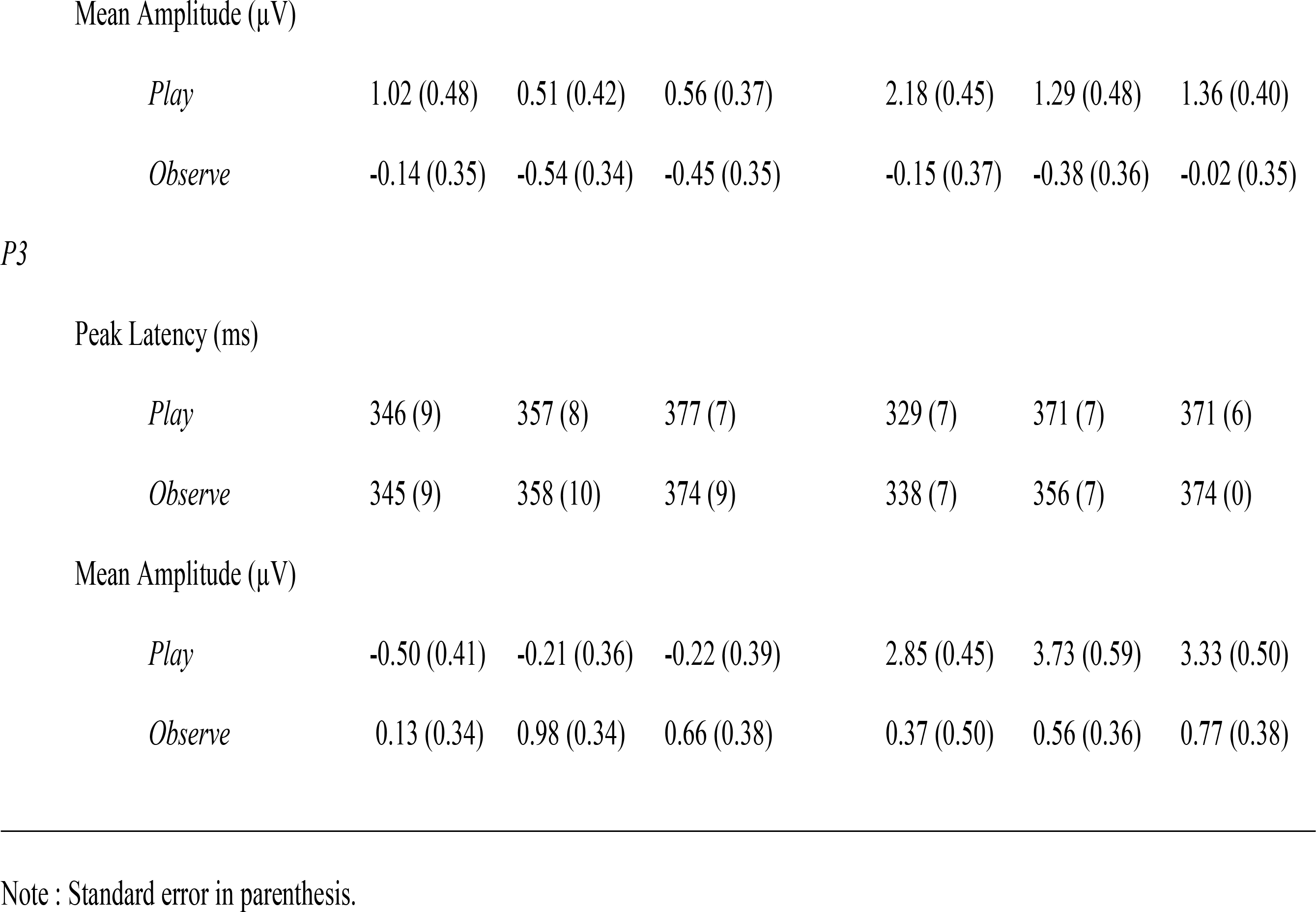
Descriptive statistics for N1, FRN and P3 components in Experiment 2

|  | *Random Opponent* |  |  | *Strategic Opponent* |  |  |
| --- | --- | --- | --- | --- | --- | --- |
|  | *Win* | *Lose* | *Draw* | *Win* | *Lose* | *Draw* |
| *N1* |  |  |  |  |  |  |
| Peak Latency (ms) |  |  |  |  |  |  |
| *Play* | 185 (2) | 182 (3) | 179 (3) | 179 (3) | 178 (3) | 178 (3) |
| *Observe* | 180 (3) | 178 (3) | 178 (4) | 177 (3) | 178 (3) | 175 (3) |
| Mean Amplitude (μV) | | | | | | |
| *Play* | -0.50 (0.41) | -0.21 (0.36) | -0.22 (0.39) | -1.14 (0.43) | -0.61 (0.40) | -0.61 (0.43) |
| *Observe* | 0.13 (0.35) | 0.98 (0.34) | 0.66 (0.39) | 0.10 (0.35) | 0.47 (0.39) | 0.51 (0.43) |
| *FRN* |  |  |  |  |  |  |
| Peak Latency (ms) |  |  |  |  |  |  |
| *Play* | 282 (5) | 282 (4) | 279 (4) | 284 (5) | 274 (4) | 268 (5) |
| *Observe* | 288 (4) | 286 (5) | 283 (5) | 288 (5) | 288 (5) | 284 (5) |
| Mean Amplitude (μV) | | | | | | |
| *Play* | 1.02 (0.48) | 0.51 (0.42) | 0.56 (0.37) | 2.18 (0.45) | 1.29 (0.48) | 1.36 (0.40) |
| *Observe* | -0.14 (0.35) | -0.54 (0.34) | -0.45 (0.35) | -0.15 (0.37) | -0.38 (0.36) | -0.02 (0.35) |
| *P3* |  |  |  |  |  |  |
| Peak Latency (ms) |  |  |  |  |  |  |
| *Play* | 346 (9) | 357 (8) | 377 (7) | 329 (7) | 371 (7) | 371 (6) |
| *Observe* | 345 (9) | 358 (10) | 374 (9) | 338 (7) | 356 (7) | 374 (0) |
| Mean Amplitude (μV) | | | | | | |
| *Play* | -0.50 (0.41) | -0.21 (0.36) | -0.22 (0.39) | 2.85 (0.45) | 3.73 (0.59) | 3.33 (0.50) |
| *Observe* | 0.13 (0.34) | 0.98 (0.34) | 0.66 (0.38) | 0.37 (0.50) | 0.56 (0.36) | 0.77 (0.38) |

Note : Standard error in parenthesis.

**Table 4.**
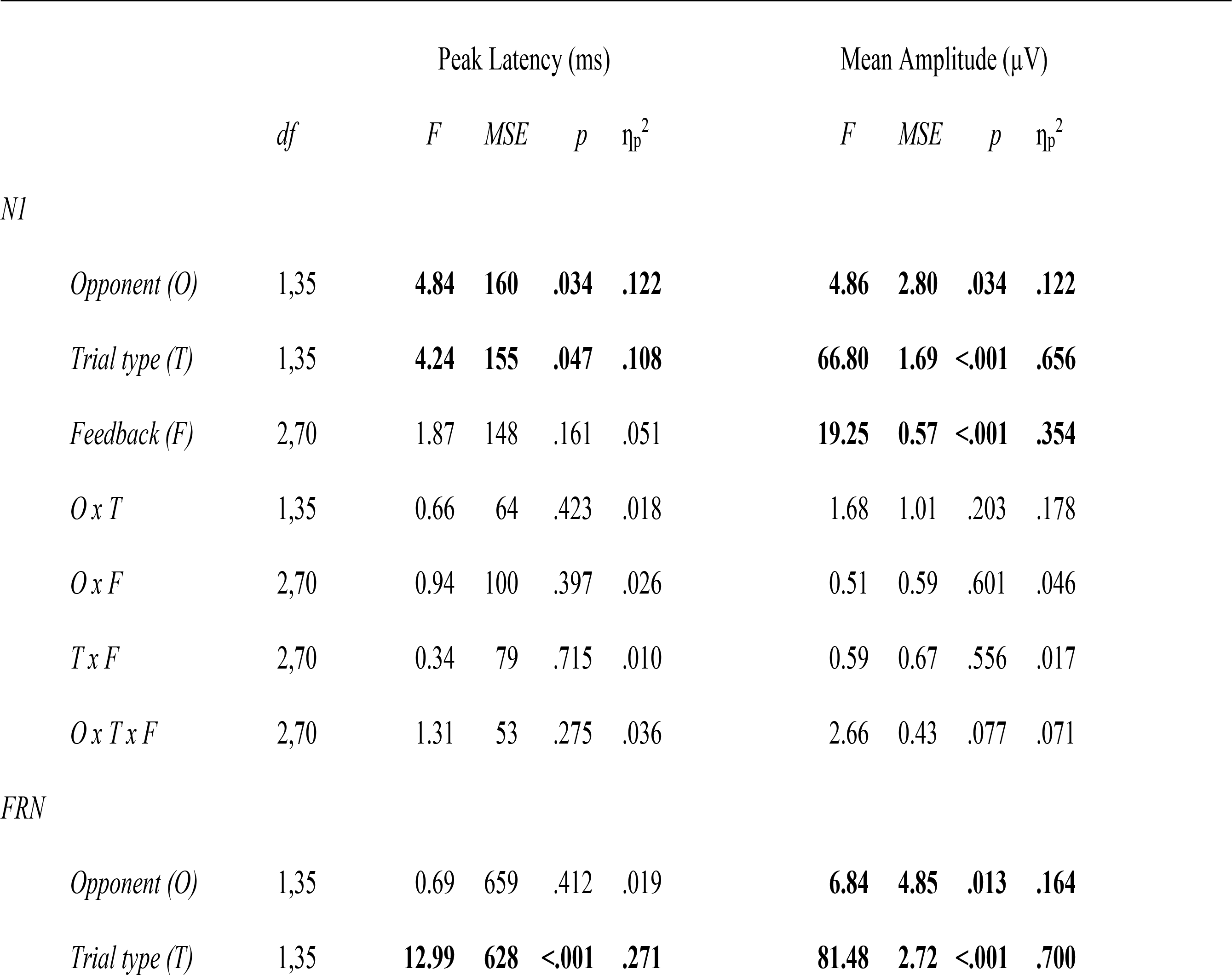
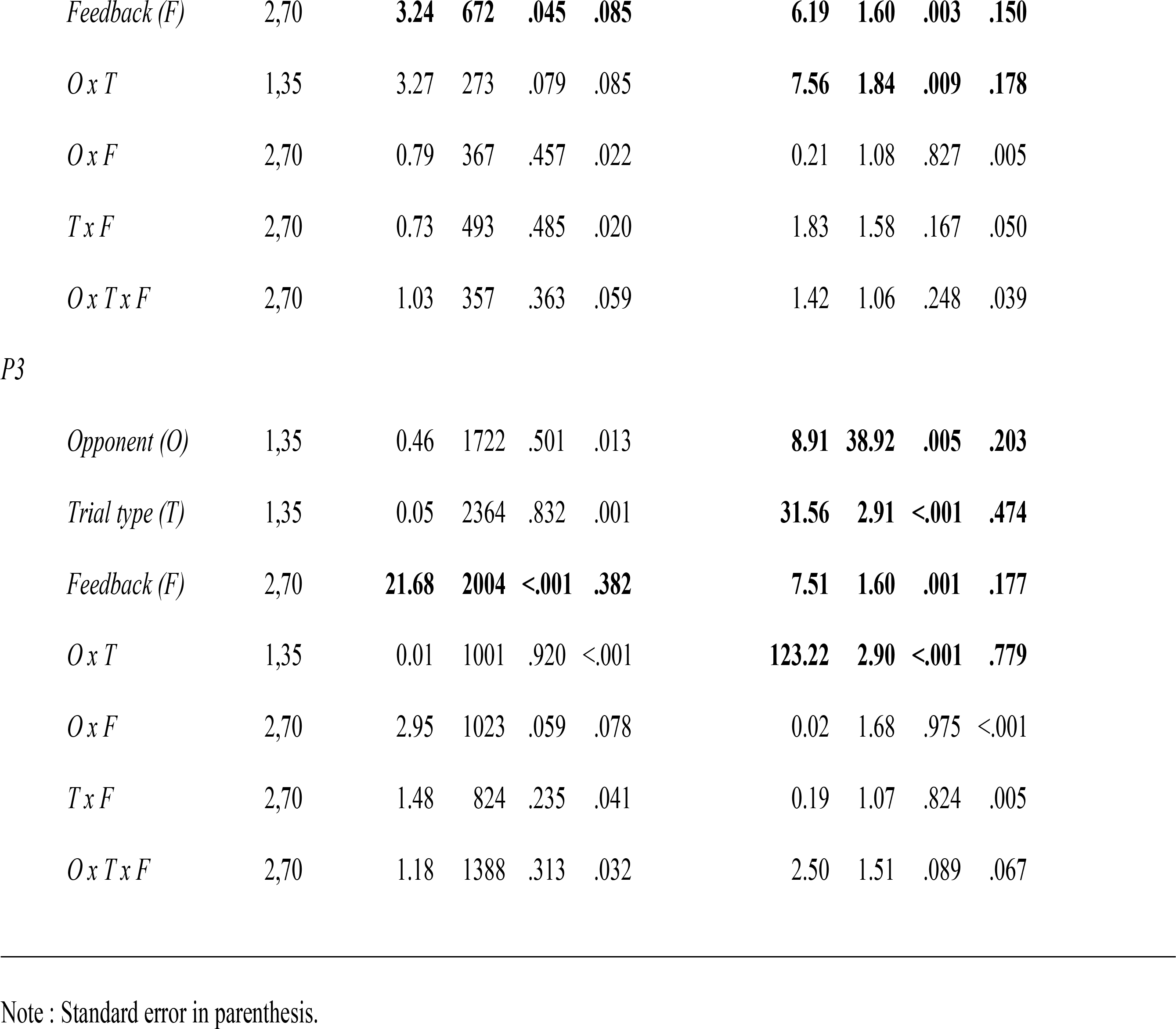
Inferential statistics for N1, FRN and P3 peak latency and mean amplitude in Experiment 2

|  |  | Peak Latency (ms) |  |  |  | Mean Amplitude (μV) |  |  |  |
| --- | --- | --- | --- | --- | --- | --- | --- | --- | --- |
|  | *df* | *F* | *MSE* | *p* | η<sub>p</sub><sup>2</sup> | *F* | *MSE* | *p* | η<sub>p</sub><sup>2</sup> |
| *N1* |  |  |  |  |  |  |  |  |  |
| *Opponent (O)* | 1,35 | **4.84** | **160** | **.034** | **.122** | **4.86** | **2.80** | **.034** | **.122** |
| *Trial type (T)* | 1,35 | **4.24** | **155** | **.047** | **.108** | **66.80** | **1.69** | **<.001** | **.656** |
| *Feedback (F)* | 2,70 | 1.87 | 148 | .161 | .051 | **19.25** | **0.57** | **<.001** | **.354** |
| *O x T* | 1,35 | 0.66 | 64 | .423 | .018 | 1.68 | 1.01 | .203 | .178 |
| *O x F* | 2,70 | 0.94 | 100 | .397 | .026 | 0.51 | 0.59 | .601 | .046 |
| *T x F* | 2,70 | 0.34 | 79 | .715 | .010 | 0.59 | 0.67 | .556 | .017 |
| *O x T x F* | 2,70 | 1.31 | 53 | .275 | .036 | 2.66 | 0.43 | .077 | .071 |
| *FRN* |  |  |  |  |  |  |  |  |  |
| *Opponent (O)* | 1,35 | 0.69 | 659 | .412 | .019 | **6.84** | **4.85** | **.013** | **.164** |
| *Trial type (T)* | 1,35 | **12.99** | **628** | **<.001** | **.271** | **81.48** | **2.72** | **<.001** | **.700** |
| *Feedback (F)* | 2,70 | **3.24** | **672** | **.045** | **.085** | **6.19** | **1.60** | **.003** | **.150** |
| *O x T* | 1,35 | 3.27 | 273 | .079 | .085 | **7.56** | **1.84** | **.009** | **.178** |
| *O x F* | 2,70 | 0.79 | 367 | .457 | .022 | 0.21 | 1.08 | .827 | .005 |
| *T x F* | 2,70 | 0.73 | 493 | .485 | .020 | 1.83 | 1.58 | .167 | .050 |
| *O x T x F* | 2,70 | 1.03 | 357 | .363 | .059 | 1.42 | 1.06 | .248 | .039 |
| *P3* |  |  |  |  |  |  |  |  |  |
| *Opponent (O)* | 1,35 | 0.46 | 1722 | .501 | .013 | **8.91** | **38.92** | **.005** | **.203** |
| *Trial type (T)* | 1,35 | 0.05 | 2364 | .832 | .001 | **31.56** | **2.91** | **<.001** | **.474** |
| *Feedback (F)* | 2,70 | **21.68** | **2004** | **<.001** | **.382** | **7.51** | **1.60** | **.001** | **.177** |
| *O x T* | 1,35 | 0.01 | 1001 | .920 | <.001 | **123.22** | **2.90** | **<.001** | **.779** |
| *O x F* | 2,70 | 2.95 | 1023 | .059 | .078 | 0.02 | 1.68 | .975 | <.001 |
| *T x F* | 2,70 | 1.48 | 824 | .235 | .041 | 0.19 | 1.07 | .824 | .005 |
| *O x T x F* | 2,70 | 1.18 | 1388 | .313 | .032 | 2.50 | 1.51 | .089 | .067 |

Note : Standard error in parenthesis.

**Figure 5.**
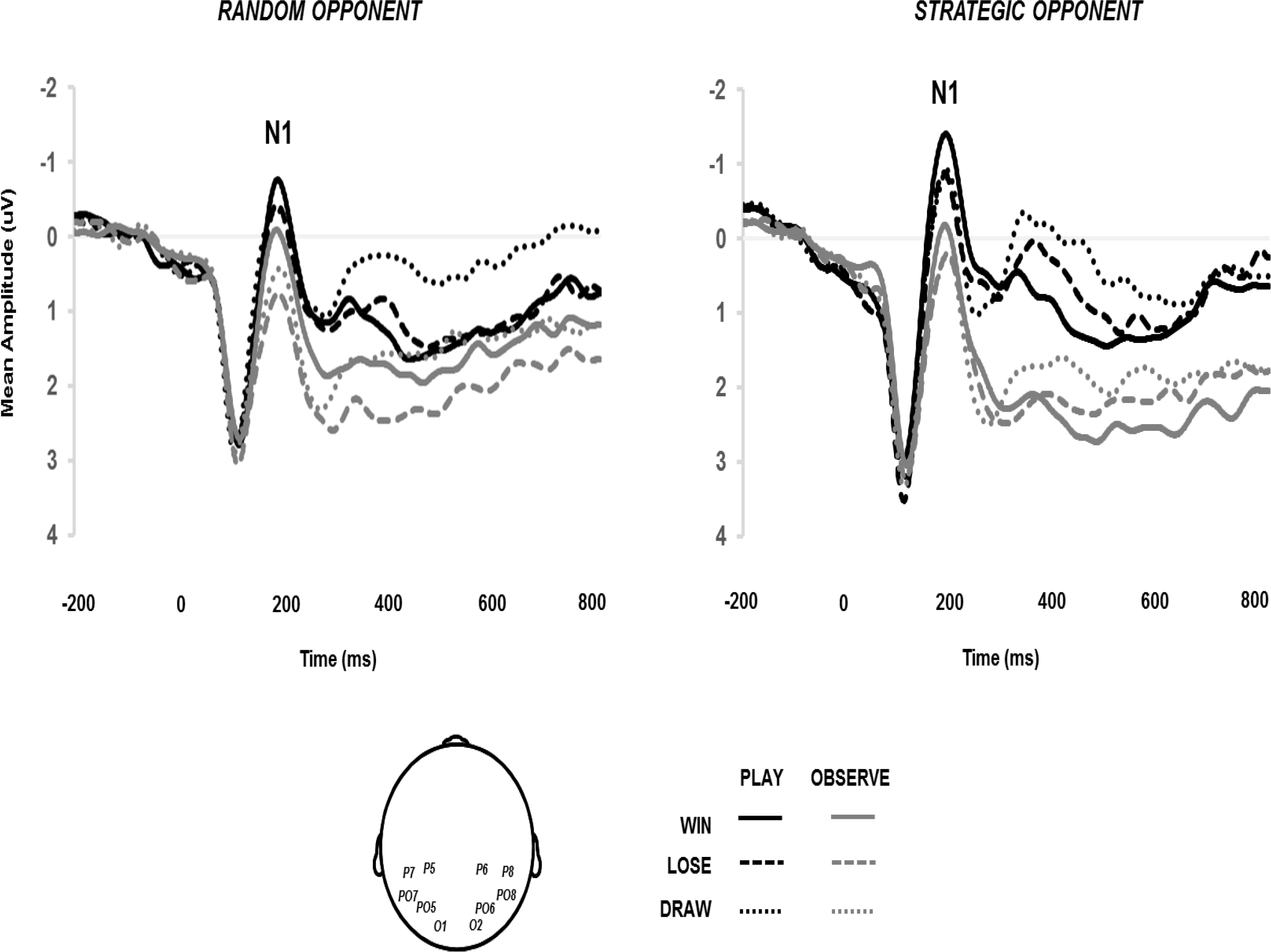
Group-average ERP from 10 parietal-occipital sites generated by the presentation of trial feedback (*win, lose, draw*) and according to the nature of the trial (*play, observe*) and opponency (*random*, *strategic*) in Experiment 2 (20 Hz filter applied for presentation).

#### Feedback-related negativity (FRN)

FRN peak latency analysis was analysed in an identical way to N1 (see Tables 3 and 4, and, Figure 6). A main effect of trial type (*p* < .001) revealed speeded FRN peak latency for *play* trials relative to *observe* trials (277 and 286 ms, respectively), and a main effect of feedback (*p* = .045) revealed speeded FRN peak latency for *draw* trials relative to *win* and *lose* trials (278, 285 and 282 ms, respectively; statistically significant for the *draw* – *win* comparison only, Tukey’s HSD *p* < .05). The FRN speeding associated with *play* trials and *draw* trials again replicated Experiment 1. Mean amplitude yielded main effects of opponent (*p* = .013), trial type (*p* < .001) and feedback (*p* = .003), in addition to a two-way interaction between opponent x trial type (*p* = .009). The data replicated the standard finding of less negative FRN for *wins* (0.73 µV) relative to both *losses* (0.22 µV) and *draws* (0.36 µV; Tukey’s HSD, *p* <.05). The interaction between opponent x trial type showed all differences between *random play* (0.70 µV), *random observe* (−0.38 µV), *strategic play* (1.61 µV) and *strategic observe* (−0.18 µV) conditions to be significant, apart from between *random observe* and *strategic observe* contexts.

**Figure 6.**
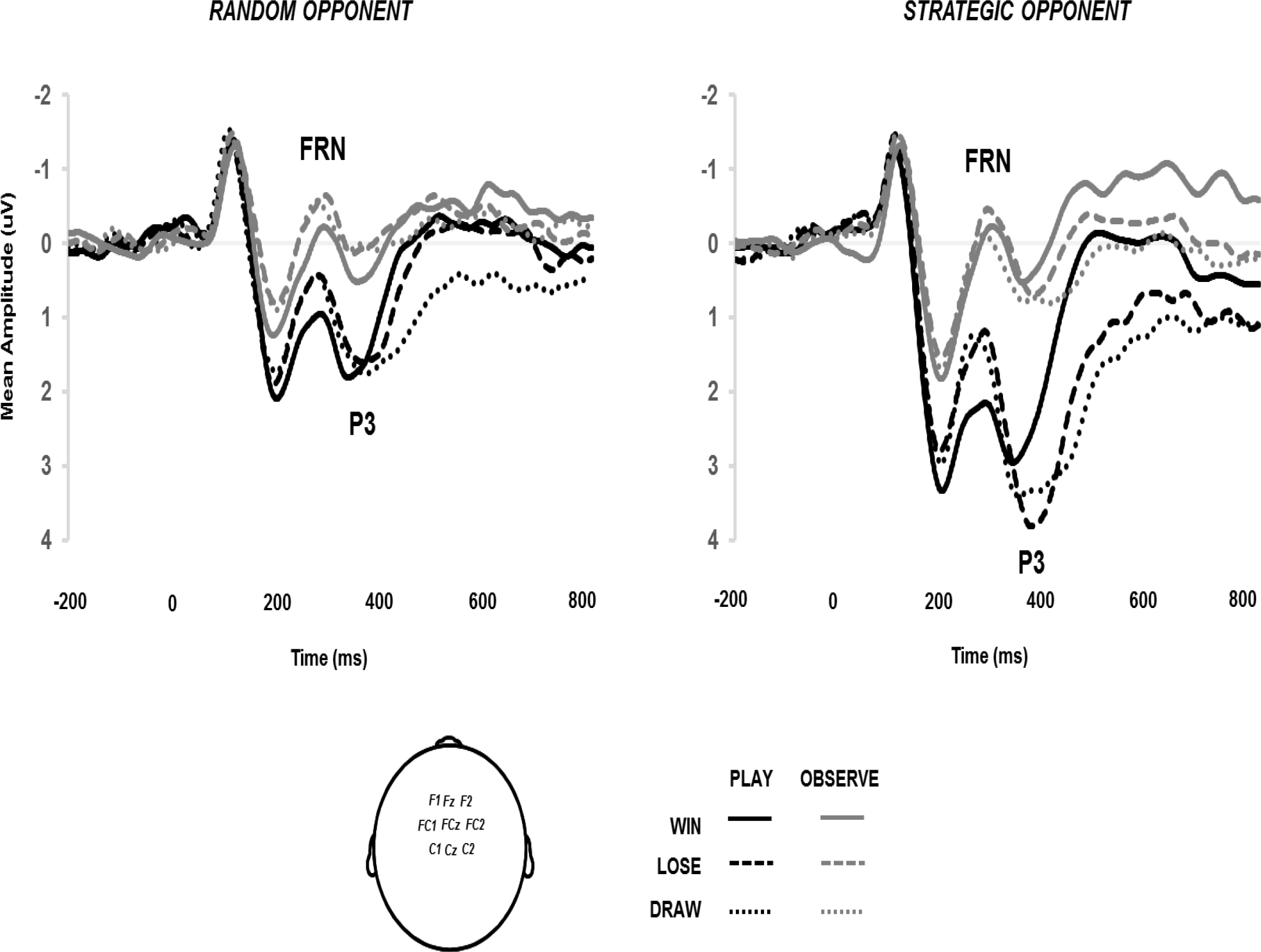
Group-average ERP from 9 fronto-central sites generated by the presentation of trial feedback (*win, lose, draw*) and according to the nature of the trial (*play, observe*) and opponency (*random*, *strategic*) in Experiment 2 (20 Hz filter applied for presentation).

The contribution of visual attention on FRN was assessed in the same way as Experiment 1, yielding a significant positive correlation during *random* opponency (*r* = .398; *p* = .016; replicating Experiment 1, see Figure 4A) but no significant correlation during *strategic* opponency (*r* = .168; *p* = .327). Empathy self-report was not correlated with FRN difference generated during *random* opponency (*r* = .174, *p* = .311), failing to replicate Experiment 1 (see Figure 4B)^1^.

#### P3

Our examination of P3 across the mid-line revealed evidence of the appropriate capture of P3a modulation at frontal, fronto-central and central sites, in the absence of any distinction P3b modulation at centro-parietal and parietal sites (see Supplementary Graph A). As a result, P3a will serve as the focus of analysis. P3 peak latency analysis returned a main effect of feedback (*p* < .001) showing an earlier peak for *win* trials relative to both *lose* and *draw* trials (340, 360 and 374 ms, respectively; Tukey’s HSD, *p* < .05). Mean amplitude revealed main effects for opponent (*p* = .005), trial type (*p* < .001) and feedback (*p* = .001), in addition to a two-way interaction between opponent x trial type (*p* < .001). P3 mean amplitude was significantly smaller for *wins* trials relative to both *lose* and *draw* trials (0.71 µV, 1.26 µV and 1.13 µV, respectively; Tukey’s HSD, *p* < .05). As per FRN mean amplitude, the opponent x trial type interaction showed that all P3 differences between *random play* (−0.31 µV), *random observe* (0.59 µV), *strategic play* (3.30 µV) and *strategic observe* (0.56 µV) conditions to be significant, apart from the comparison between *random observe* and *strategic observe* contexts. P3 was clearly largest during *strategic play* for all of the three possible trial results (*win, lose, draw*).

To assess the contribution of outcome frequency on P3 amplitude, P3 differences between *lose* and *win* during the *strategic play* condition were correlated with the difference between win and lose rates generated during the *strategic play* condition for all 36 participants. The positive correlation between P3 mean amplitude difference between *lose* and *win*, and, *win* – *lose* rate difference was significant (*r* = .355; *p* = .034). This suggests that as wins become more frequent and losses become more rare as a function of successful learning, the larger the P3 difference between losses and wins becomes.

To assess the overall contribution of FRN on P3, FRN differences between *lose* and *win* were calculated alongside P3 differences between *lose* and *win*, for each of the four conditions in the study. The correlation between these four data points was then calculated on an individual basis, and compared at a group level to the test statistic of 0. With an average correlation of .26 (SE = .09) and *t*(35) = 2.88, *p* = .006, there was evidence to suggest that the magnitude of difference between *wins* and *losses* reflected in the FRN was similarly reflected in the later P3 (see also Zhou et al., 2010).

## Discussion

Experiment 2 revealed how the removal of response choice impacted on competitive performance against both *random* and *strategic* opponents at a phasic (rather than tonic; Experiment 1) level. The behavioural data demonstrated that as a group there was some learning of optimal play against the exploitable (*strategic*) opponent, and that the ability to exhibit this behaviour was modulated by both previous trial type and previous outcome. Firstly, the removal of response choice in the preceding trial (*observe* trial *n-1*) interrupted participants’ ability to reengage optimal play on the current play trial *n*, as indexed by a reduction in correct strategy implementation for *observe-play* trials relative to *play-play* trials. Therefore, in the context of *strategic* opponency, the removal of response choice had an inhibitory effect on performance. Secondly and in alignment with previous results, participants were more likely to engage in higher-quality decision making following a positive outcome (*win*) relative to a negative outcome (*lose* or *draw;* Dyson et al., 2016; Forder & Dyson, 2016; Laakasuo et al., 2015; Mitzenmacher & Upfal, 2005). This was despite the fact that the strategy itself was a form of *shift* behaviour. Therefore, the reduced ability to initiate *shift* behaviour following negative outcome and the enhanced ability to initiate *shift* behaviour following positive outcome was in contrast to the very basic reinforcement learning tendencies of *lose-shift*, and, *win-stay*.

We failed to find evidence for our secondary hypothesis that in the context of *random* opponency, the removal of response choice should have had a facilitatory effect on performance in moving players towards MES. While the data from Experiment 2 suggest a somewhat pessimistic (inhibitory) role of interruption during competition, it is clear that similar kinds of decision-making might be naturally interrupted after the experience of a positive outcome either by the player in the form of post-reinforcement pausing (e.g., Dixon & Schreiber, 2004; Dyson et al., 2018; Verbruggen et al., 2017; Forder & Dyson, 2016) or by the opponent itself such as in the case of slot machines where longer music tends to play when the win is bigger (e.g., Dixon et al., 2013). This leads us to the intriguing possibility that mandatory pauses following negative outcomes during *play* conditions might help to break the cyclical poorer-quality decision making characterised in problem gambling (see Ivan, Banks, Goodfellow & Gruber, 2018, for a similar suggestion).

The neural data of Experiment 2 replicate and extend much of Experiment 1 in terms of N1 and FRN ^1^. In particular, against unexploitable opponents, the N1 significantly contributed to the extent to which negative and positive outcomes were perceived as distinct categories by the further down-stream FRN. However, the added value of Experiment 2 was demonstrating the modulation of P3 by both outcome frequency (Zheng et al., 2017) and the motivational significance of certain classes of trial (Nieuwenhuis et al., 2005; Polich, 2003). First, we confirmed that individuals exhibiting high win rates and low loss rates against exploitable (*strategic*) opponents also showing larger differences between losses and wins in P3 amplitude. Second, P3 amplitude was maximal for *play* trials against a *strategic* opponent, and for negative relative to positive outcomes. It is worth being explicit here that the four combinations of different trial types (*random play*, *random observe*, *strategic play*, *strategic observe*) were equi-probable across the study. Thus, P3 modulation in response to trial type must be a function of motivational significance and cannot be a function of frequency. The idea that P3 reacts the strongest to unexpected, rare negative outcomes in the context of potential exploitation of a strategic opponent is critical for the view that P3 acts as an intermediary between the initial FRN reaction to outcome on the current trial and behavioural performance on the subsequent trial.

## General Discussion

The current studies sought to examine the behavioural and neural dynamics of competitive decision making as a function of the positive or negative nature of trial outcome, whether an individual was more or less responsible for the outcome (i.e., *play* versus *observe* trials), and, whether or not there was vulnerability in the opponent’s behaviour (i.e., *strategic* versus *random* opponents).

At a behavioural level, the data were especially instructive in revealing the limitations of some basic reinforcement learning tendencies. Against unexploitable opponents, both Experiment 1 and 2 showed no evidence of *win-stay* behaviour and achieved a rough approximation of mixed-equilibrium strategy (MES) following positive outcomes (see also Dyson et al., 2016). This is not to say that *win-stay* behaviour never occurs in these contexts, and indeed we have evidence that action repetition following reward can be made more frequent by increasing the value of reward (Forder & Dyson, 2016), or, being explicit that the opponent from the previous trial has returned for the next trial (Srihaput, Sundvall & Dyson, in preparation). Rather, the expression of *win-stay* appear to be somewhat less rigid than the expression of *lose-shift*, consistent with traditional idea that losses are more important to the organism than wins (Kahneman & Tversky, 1979; although see Yechlam, 2018, for an alternative view).

These findings are notable as much has been made both historically and contemporarily of the so-called ‘bounded rationality’ of the individual (e.g., Simon, 1956; Griessinger & Coricello, 2015), the degree to which individuals are beholden to reinforcement learning principles (Skinner, 1957), and, the difficulties humans have in exhibiting random behaviour (Neuringer, 1986; Terhune & Brugger, 2011). Challenges to the putatively ingrained nature of reinforcement learning rules such as *win-stay*, *lose-shift* are further strengthened by the behaviour observed in both *strategic* and *random* conditions in Experiment 2. Recall that in the *strategic* condition, successful performance was defined as the increased use of a *downgrade* strategy, which guaranteed a winning outcome on 80% of *play* trials. Note that this particular exploitation strategy represents a form of *shift* behaviour, historically promoted by exposure to negative rather than positive outcomes (i.e., *lose-shift*). The fact that *shift* behaviour was at its highest following *win* trials in Experiment 2 highlights that individuals can overcome *win-stay* tendencies and adopt *win-shift* behaviours when the environment requires it (similar to the behaviour of nectarivorous birds and species that forage in environments with a fast depletion rate; Stagner, Michler, Rayburn-Reeves, Laude & Zentall, 2013). Even more dramatic deviation from reinforcement principles was observed against a *random* opponent, where participants approximated MES strategy across both positive (*win*) and negative (*lose*, *draw*) outcomes (contra Neuringer, 1986; Terhune & Brugger, 2011). While we might have anticipated variation in the more malleable *win-stay* behaviour, the reduction of *lose-shift* behaviour to almost MES performance reminds us that the behavioural tendencies following both reward and punishment can be controlled. It is important for future research to specify under what conditions we believe outcome-independent, random behaviour to be facilitated. It would seem to us that when participants experience a *relative* and *discrete* comparison between different styles of opponent (e.g., *random* and *strategic*) within the same experimental session, this improves the likelihood of observing random behaviour against unexploitable opponents. Our lab has since replicated this finding in a separate study (Sundvall & Dyson, in preparation) where the critical design feature again appear to be the comparison between *random* and *strategic* opponents in separate blocks.

The neural data recorded concurrently during the session also revealed a number of replicable findings that further speak to the dynamics of competitive decision-making and performance. In both Experiments 1 and 2, we found that the visual attention allocated to *observe* trials was reduced relative to *play* trials, as indexed by smaller N1 amplitude. This was the case both when response selection was removed at a tonic (block; Experiment 1) and phasic (trial-to-trial; Experiment 2) level. The smaller N1 during *observe* trials also likely gave rise to the finding that FRN amplitudes were also generally more negative during *observe* trials. This is because the N1 deflection at parieto-occipital sites (e.g., Figure 1a) reverses in polarity at fronto-central sites (e.g., Figure 1b) as reflected in the positive-going deflection around 200 ms after outcome on-set in Figure 2B (see also Bellebaum & Daum, 2008; Osinsky, Mussel & Hewig, 2012). Indeed the close relationship between N1 and FRN was further supported by the positive correlation between *play* – *observe* mean amplitude differences during *random* opponency shown in Figure 3A. Therefore, the degree of visual attention allocated to individual trials indexed by N1 determined the degree to which wins and losses were neurally registered as distinct categories of outcome by FRN. Furthermore, the neural distinction between trials for which one was (*play*) and was not (*observe*) responsible for outcome was more pronounced at the tonic (Experiment 1) rather than phasic (Experiment 2) level, as supported by a between-experiment interaction with trial type for FRN amplitudes during *random* opponency [F(1,70) = 13.06, MSE = 2.82, *p* < .001, ƞ_p_^2^ = .157]. It is worth noting that in terms of the top-down control that individuals can bring to bear upon the FRN signature, FRN amplitudes during *play* trials did not modulate across Experiments 1 and 2 (0.99 versus 0.70, respectively) but FRN amplitude during *observe* trials did (−1.25 versus −0.38, respectively). However, it is also important to note that there are broad similarities in FRN sensitivity across Experiments 1 and 2 against *random* opponency, despite the fact that quite different behaviour was expressed. In support of the former point, a between-experiment comparison only reveals a main effect for feedback [F(2,140) = 11.16, MSE = 1.27, *p* < .001, ƞ_p_^2^ = .137] without any higher-order interactions. Contrast this with the observation that in Experiment 1, *lose-shift* behaviour was in evidence while in Experiment 2 participants exhibited MES. This disconnect between brain and behaviour raises questions over the functional link between the registration of outcome by FRN and the subsequent behavioural consequences of outcome. Thus, FRN may be a necessary but not sufficient condition for causing behavioural updating in the light of previous performance (e.g., Müller, Möller, Rodriguez-Fornells & Münte, 2006).

Variation in the P3 component in Experiment 2, and its relation to the earlier FRN and later behaviour, helps to bridge this functional gap. First, we showed that P3 amplitude was modulated by outcome frequency in the context of competitive decision-making (Zheng et al., 2017). This should serve as a cautionary finding for studies that wish to interpret the P3 solely as the brain’s reward response (Fielding et al., 2018; see Dyson et al., 2018, for more details). Second, in terms of motivational significance during *observe* trials, neither P3 amplitude nor FRN amplitude was sensitive to the *random* or *strategic* nature of the opponent. Rather, it was only when participants were able to *play* with their opponent (defined here in terms of the ability to make response selections) then P3 amplitude (and FRN amplitude) became further enhanced as a function of competing against an exploitable (*strategic*) rather than unexploitable (*random*) opponent. This aligns closely with the view of Olivera et al. (2007) where the P3 serves as a gain function for motivationally salient events. In these regards, the P3 appears to be in tune with both degree of responsibility for the current outcome, and, the informational value of the current outcome in terms of mental model updating, but only when there are reliable environmental contingencies such as when an opponent is predictable and hence exploitable. More specifically, when the participant cannot choose their response for the upcoming trial, then it follows that they are less directly responsible for any outcomes that result, hence the motivational significance of an *observe* trial is lower than that of a *play* trial. Furthermore, with the eventuality that their opponent cannot be exploited in the *random* condition, there is no mental model of opponent performance worth developing that could increase the likelihood of win maximization. In contrast, outcome information is of greater informational value during the *strategic* condition, where an accurate model of opponent predictability will yield a higher win rate.

It is also worth noting that the use of the RPS paradigm in studying neural responses to outcome is beneficial, in that the examination of *draw* trials (as well as *win* and *lose* trials) satisfies the need to understand how ambiguous or neutral outcomes relate to more-clearly defined successes and failures (Holroyd et al., 2006; Müller et al., 2006; Gu, Feng, Broster, Yuan, Xu & Luo, 2017). For example, the interpretation of a *draw* as a near-win or near-loss might differ according to an individual’s stance towards gambling behaviour (Dixon et al., 2013; Ulrich & Hewig, in press). Behaviourally, the data from *draw* trials in Experiment 2 are particularly intriguing as optimal downgrade performance appears worse following *draw* feedback relative to the more negatively valanced *lose* feedback (see Figure 4). Moreover, the current observations from Experiments 1 and 2 that *draw* feedback elicit earlier visual N1 and FRN peak responses, as well as intermediate FRN mean amplitude situated between unambiguously positive (*win*) and negative (*lose* and *draw*) feedback, suggest subtle distinctions between potentially ambiguous forms of feedback that future work would do well to pursue (e.g., Gu et al., 2017). Future work should also strengthen the examination of behavioural effects in the neural domain, as questions remain regarding the neural expression of sequential effects observed in the behavioural data. For example, the phasic shifts between *play* and *observe* trials in Experiment 2 clearly impacted on behavioural output during the strategic condition. However, our examination of neural inter-trial contingencies between *play* or *observe activity* at trial *n* as a function of *play* or *observe* activity at trial *n-1* (see Supplementary Materials B) did not reveal any interaction. Continued experimentation where inter-trial contingencies are at the heart of design will help to adjudicate between the absence of effect and reduced signal-to-ratio ratios.

In sum, decision-making during competition often requires high-quality thought and action for success but often yields poor-quality thought and action as a result of the charged nature of the environment. Appreciating the conditions under which we may control the quality of decision-making to see it rise, and the conditions under which we cannot control the quality of decision-making only to see it fall, is critical for understanding how we approach, engage and ultimately remove ourselves from increasingly sophisticated competitive environments. The continued value of studying deceptively simple games is that they allow high degree of control and yield clear guidelines for the mechanisms in helping individuals improve the quality of decision-making when they may naturally be at their most cognitively vulnerable.

## Supporting information

Data Experiment 2

Data Experiment 1

## Acknowledgements

The work was partly supported by a Research Development Fund from the University of Sussex under grant SA016-01 awarded to BJD. Data from Experiment 1 was previously presented at The Society for Applied Research in Memory and Cognition, 3-6th January 2017, Sydney, Australia. Some of the ERP data from Experiment 2 appeared for illustrative purposes only in Dyson, Forder & Sundvall (2018). Correspondence should be addressed to: Ben Dyson, Department Of Psychology P-217 Biological Sciences Building, University of Alberta, Edmonton, AB, T6G 2E9, Canada.

## Footnotes

1 On the basis of our previous data (Forder & Dyson, 2016) and given the inability to build a reliable mental model of opponent performance with an unexploitable opponent, we had not anticipated observing reliable P3 activity in Experiment 1. However, in light of the results from Experiment 2 and at the request of reviewers, the same time windows for peak latency and mean amplitude measurements used in Experiment 2 were used to identify P3a activity in Experiment 1, where participants only interacted with a random opponent under conditions of *play* and *observation*. Similar to Experiment 2, peak latency measurements revealed a main effect of feedback (F[2,70] = 10.50, MSE = 1597, *p* < .001, ƞ_p_^2^ = .231) wherein *draw* latencies were significantly slower than both *lose* and *win* outcomes (356, 336, 326 ms, respectively; Tukey’s HSD, *p* < .05). Trial type main effect (F[1,35] = 2.99, MSE = 3104, *p* = .093, ƞ_p_^2^ = .079) and trial type x feedback interaction (F[2,70] = 0.27, MSE = 1385, *p* = .768, ƞ_p_^2^ = .008) were not significant. Mean amplitude analysis revealed a main effect of trial type (F[1,35] = 79.57, MSE = 5.51, *p* < .001, ƞ_p_^2^ = .694) similar to the relevant data from Experiment 2 in that *random play* conditions yielded greater positivity (1.67 µV) than *random observe* conditions (−1.18 µV). This was in the absence of a main effect of feedback (F[2,70] = 2.65, MSE = 1.36, *p* = .078, ƞ_p_^2^ = .070) and interaction (F[2,70] = 0.26, MSE = 0.89, *p* = .261, ƞ_p_^2^= .007). An examination of ERPs down the mid-line suggested P3b activity in alignment with FRN activity, where *wins* continued to be less negative (more positive) than *losses* or *draws* (see Supplementary Graph A2).

## Author Contributions

BJD designed and wrote the experimental scripts for Experiments 1 and 2. LF collected the data and carried out initial data analyses for Experiment 1. BAS and TM collected the data and carried out initial data analyses for Experiment 2. BJD finalized data analyses and wrote the first draft of the manuscript. BAS, TM and LF commented on the manuscript.

