## Supplementary A for "Behavioural and neural limits in competitive decision making: The roles of outcome, opponency and observation"

*Supplementary Materials*

First, engagement with specific blocks of RPS was assessed using a slightly-modified Game Engagement Questionnaire (GEQ; Brockmyer, Fox, Curtiss, McBroom, Burkhart & Pidruzny, 2009). 18 from 19 original items were evaluated on a 5 point scale, encompassing the factors of *absorption*, *flow*, *presence* and *immersion*. The 1 item to be dropped from the revised GEQ was ‘I play longer than I meant to’ (related to *presence*). This item was deemed inappropriate given the fixed number of rounds in all RPS block. All items were also re-written to refer to the past tense (e.g., ‘I lose track of time’ became ‘I lost track of time’). Second, the degree of anthropomorphism attributed to the computer opponent was assessed on the basis of Epley, Akalis, Waytz & Cacioppo (2008, Study 1) where five anthropomorphic states (‘mind of its own’, ‘intentions’, ‘free will’, ‘consciousness’, ‘experienced emotion’) and three non-anthropomorphic states (‘attractive’, ‘efficient’, ‘strong’) were adjudicated on an 11-point scale (c.f,. Waytz, Cacioppo & Epley, 2010, Box A1). Third, aspects of self-reported co-presence and perceived other’s co-presence were adapted from the scales provided by Nowak & Biocca (2003). 7 modified items, evaluated on a 5 point scale, were deemed appropriate for opponent interaction in the context of RPS (3 related to self-reported co-presence, 4 related to perceived other’s co-presence; see Appendix). A further 5 unique items were added to this particular questionnaire to assess the participant’s understanding of the opponent’s strategy: “I felt as though my opponent had a strategy that was based on the moves I was making” (e.g., *other* strategy), “I felt as though my opponent had a strategy that was based on the moves it was making” (e.g., *self* strategy), “My opponent exhibited a human-like strategy”, “I felt like my opponent was somehow cheating” and “I found this block of RPS rewarding to play.” Finally, to assess the empathy of the participant, the Toronto Empathy Questionniare (TEQ; Spreng, McKinnon, Mar & Levine, 2009) was administered after all blocks of RSP have been played. The TEQ consists of 16-items (in contrast to the 80-item Empathy Quotient; Baron-Cohen & Wheelwright, 2004) judged on a 5 point scale, yielding a single empathy factor.

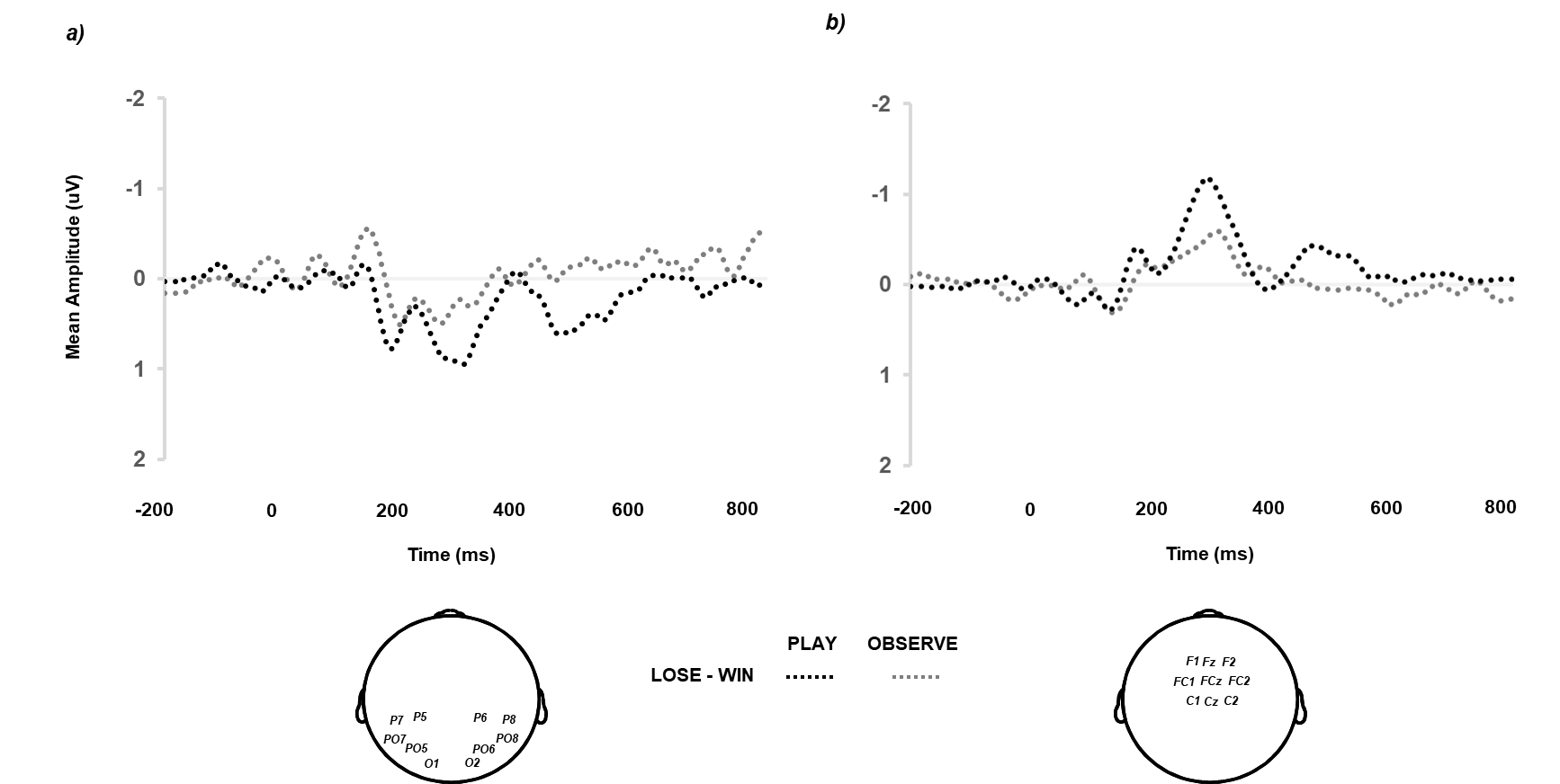

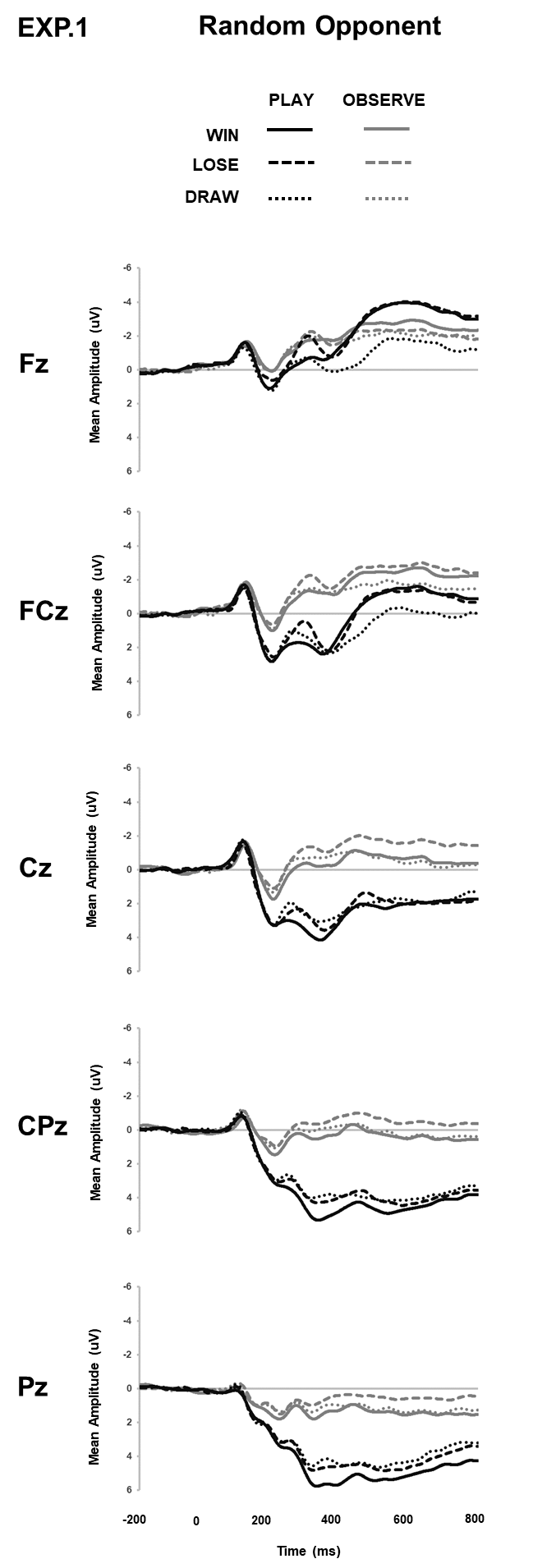

Figure Caption

Supplementary Figure A1. Group-average ERP from a) 10 parieto-occipital and b) 9 fronto-central sites generated by the difference between lose and win trials as a function of trial type (play, observe; 20 Hz filter applied for presentation).

Supplementary Figure A2. Group-average ERP generated by the presentation of trial feedback (win, lose, draw) as a function of trial type (play, observe) across 5 mid-line electrodes from anterior (Fz) to posterior (Pz) sites (20 Hz filter applied for presentation).

*References*

Baron-Cohen, S. & Wheelwright, S. (2004). The empathy quotient: An investigation of adults with Asperger syndrome or high functioning autism, and normal sex differences. *Journal of Autism and Developmental Disorders*, *34*, 163-175.

Brockmyer, J. H., Fox, C. M., Curtiss, K. A., McBroom, E., Burkhart, K. M. & Pidruzny, J. N. (2009). The development of the Game Engagement Questionnaire: A measure of engagement in video game-playing. *Journal of Experimental Social Psychology*, *4*5, 624-634.

Epley, N., Akalis, S., Waytz, A. & Cacioppo, J. T. (2008). Creating social connection through inferential reproduction. *Psychological Science*, *19*, 114-120.

Nowak, K. L. & Biocca, F. (2003). The effect of the agency and anthropomorphism of users’ sense of telepresence, copresence, and social presence in virtual environments. *Presence: Teleoperators and Virtual Environments*, *12*, 481-494.

Spreng, R. N., McKinnon, M. C., Mar, R. A. & Levine, B. (2009). The Toronto Empathy Questionnaire: Scale development and initial validation of a factor-analytic solution to multiple empathy measures. *Journal of Personality Assessment*, *91*, 62-71.

Waytz, A., Epley, N. & Cacioppo, J. T. (2010). Social cognitive unbound: Insights into anthropomorphism and dehumanization. *Current Directions in Psychological Science*, *19*, 58-62.
