## Supplementary B for "Behavioural and neural limits in competitive decision making: The roles of outcome, opponency and observation"

*Behavioural Data*

*Questionnaire data*

After all four blocks of RPS had been completed participants answered four questionnaires. To assess empathy they completed the Toronto Empathy Questionnaire (TEQ; Spreng, McKinnon, Mar, & Levine, 2009) and the Interpersonal Reactivity Index (IRI; Davis, 1980). The IRI was further divided into Perspective Taking (PT), Fantasy Scale (FS), Empathic Concern (EC) and Personal Distress (PD) (Davis, 1980). The influence of PD scores on ERP responses was predicted on the basis of previous data (Larson, Fair, Good, & Baldwin, 2010). Both Sensitivity to Loss (SL; Palomäki, Laakasao, & Salmela, 2012) and Reward Responsiveness (RR; Van den Berg, Franken, & Muris, 2010) were also completed to assess participants’ relationship to competitive environments relative to their neural responses.

TEQ and IRI scores were positively correlated (*r* = .601, *p* < .001). FRN difference between *play* and *observe* conditions [(*play* *lose* – *play* *win*) – (*observe* *lose* – *observe win*)] for the *random* opponent condition was correlated with TEQ for all 36 participants. The current data (*r* = -.303, *p* = .073) failed to show significance but was consistent with the negative correlation observed by Dyson & Forder (in review). The same was not true of the correlation between FRN difference and IRI (*r* = .025, *p* = .885), nor of the PD subscale of IRI (*r* = .025, *p* = .884). Regarding the two remaining measures, SL showed a positive trend (*r* = .306, *p* = .070) and RR did not (*r* = .013, *p* = .941).

*Reaction time*

Median RT was calculated per participant for each of the three possible feedback displays (win, lose, draw) for play *n* to play *n+1* and observe *n* to play *n+1* trials, across *random* and *strategic* opponent conditions (see Table 1). 13 participants had to be rejected as a result of their average median RT being at least twice as large as the group average median RT, within any ANOVA cell (after Dyson, Sundvall, Forder & Douglas, in press). For the remaining 23 participants, a three-way repeated-measures ANOVA using opponent (*random, strategic*) trial *n* status (*play, observe*) and feedback (*win, lose, draw*) revealed main effects of opponent [F(1,22) = 12.68, MSE = 2264868, *p* = .002, ƞ_p_^2^ = .366] and trial *n* status only [F(1,22) = 28.32, MSE = 59264, *p* < .001, ƞ_p_^2^ = .563]. The main effect of feedback [F(2,44) = 0.94, MSE = 162032, *p* = .399, ƞ_p_^2^ = .041] and interactions between opponent x trial *n* status [F(1,22) = 2.67, MSE = 55602, *p* = .116, ƞ_p_^2^ = .108], opponent x feedback [F(2,44) = 0.76, MSE = 146134, *p* = .472, ƞ_p_^2^ = .034], trial *n* status x feedback [F(2,44) = 0.68, MSE = 86395, *p* = .514, ƞ_p_^2^ = .030], and, opponent x trial *n* status x feedback [F(2,44) = 3.05, MSE = 57168, *p* = .058, ƞ_p_^2^ = .122] all failed to reach significance. In these regards, the RT data were clear cut in terms of showing slower RTs against a *strategic* relative to *random* opponent (1411 vs. 766 ms), and, slower RTs following an observation trial *n* relative to a play trial *n* (1167 vs. 1011 ms).

The resultant data failed to replicate the findings of Verbruggen, Chambers, Lawrence & McLaren (2017, or, Dyson, Sundvall, Forder & Douglas (in press), in terms significant speeding following negative outcomes during *random* opponency and significant slowing following negative outcomes during *strategic* opponency. The idea that slowing following loss in the context of an exploitable opponent should positively correlate with the degree of difference between win rate and loss rate for each individual (Dyson et al., in press) also failed to reach significance but was in the correct direction (*r* = .302, *p* = .162).

*ERP data*

Electrophysiological data was reanalyzed as a function of the relationship between the *play* or *observe* status of trial *n-1* (previous trial) and trial *n* (current trial) for both *random* and *strategic* opponent conditions, resulting in separate 2 x 2 x 2 repeated-measures ANOVA for FRN and P3, respectively (see also Supplementary Tables B1 and B2, and Figure B1).

*Feedback-related negativity (FRN)*

Peak latency analyses only revealed a main effect of trial *n* (*p* < .001) essentially replicating the observation of faster FRN peak latency for *play* relative to *observe* trials as shown in the main analysis. Similarly, mean amplitude analyses replicate the main effects of opponent (*p* = .003), trial *n* (*p* < .001) and interaction between opponent x trial *n* (*p* = .004) reported in the main analysis.

*P3*

Peak latency analyses revealed an interaction between opponent x trial *n-1* (p = .024). Here, previous *play* at trial *n-1* against a random opponent yielded significantly slower P3 peak latency (364 ms; all Tukey’s HSD, *p* < .05) at trial *n*, relative to previous *observe* at trial *n-1* against a random opponent (351 ms), and, *play* *n-1* (348 ms) and *observe* *n-1* (351 ms) against a *strategic* opponent. We cautiously suggest this effect may be due to the bottom-up nature of resolving *play* outcomes against an unexploitable opponent. P3 mean amplitude analyses again replicated the main effects of opponent (*p* < .003), trial *n* (*p* < .001) and interaction between opponent x trial *n* (*p* = .005) reported in the main analysis.

Supplementary Table B1. Descriptive statistics for reaction time at trial *n* as a function of opponent, feedback at trial *n-1* and trial type at trial *n-1*

___________________________________________________________________________________________________________

*Random Opponent	Strategic Opponent*

Trial *n-1*	Trial *n*	*Win	Lose	Draw	Win	Lose	Draw*

*Play	Play*	735 (64)	683 (50)	717 (65)	1211 (181)	1422 (198)	1298 (166)

*Observe	Play*	784 (59)	853 (79)	836 (65)	1438 (183)	1482 (193)	1615 (184)

___________________________________________________________________________________________________________

Note : Standard error in parenthesis.

*Supplementary Table B2.* Descriptive statistics for FRN and P3 components for the trial *n-1* to trial *n* analyses

___________________________________________________________________________________________________________

*Random Opponent	Strategic Opponent*

*Play n	Observe n	Play n	Observe n*

*FRN*

Peak Latency (ms)

*Play n-1*	277 (4)	288 (4)	276 (4)	287 (5)

*Observe n-1*	279 (5)	282 (4)	270 (4)	283 (4)

Mean Amplitude (µV)

*Play n-1*	0.59 (0.40)	-0.36 (0.35)	1.72 (0.43)	-0.11 (0.39)

*Observe n-1*	0.88 (0.41)	-0.33 (0.31)	1.87 (0.41)	-0.10 (0.33)

*P3*

Peak Latency (ms)

*Play n-1*	359 (8)	370 (9)	344 (7)	352 (9)

*Observe n-1*	355 (9)	346 (9)	347 (6)	355 (8)

Mean Amplitude (µV)

*Play* *n-1*	1.72 (0.35)	0.24 (0.34)	2.81 (0.47)	0.62 (0.41)

*Observe* *n-1*	1.60 (0.42)	0.21 (0.31)	3.20 (0.41)	0.66 (0.29)

___________________________________________________________________________________________________________

Note : Standard error in parenthesis.

*Supplementary Table B3.* Inferential statistics for FRN and P3 components for the trial *n* to trial *n+1* analyses

___________________________________________________________________________________________________________

Peak Latency (ms)	Mean Amplitude (µV)

*df	F	MSE	p*	ƞ_p_^2^	*F	MSE	p*	ƞ_p_^2^

*FRN*

*Opponent (O)*	1,35	1.34	415	.254	.037	**10.16	2.99	.003	.225**

*Trial n-1 (Tn-1)*	1,35	3.44	229	.072	.090	1.87	0.55	.180	.051

*Trial n (Tn)*	1,35	**14.64	478	<.001	.294**	**73.05	2.19	<.001	.676**

*O x Tn-1*	1,35	0.57	361	.456	.016	0.11	0.96	.736	.003

*O x Tn*	1,35	1.68	240	.203	.046	**9.59	1.26	.004	.215**

*Tn-1 x Tn*	1,35	1.13	152	.294	.031	0.75	0.97	.393	.021

*O x Tn-1 x Tn*	1,35	1.57	274	.218	.042	0.10	0.61	.748	.003

*P3*

*Opponent (O)*	1,35	2.90	1648	.097	.076	**13.53	4.13	<.001	.279**

*Trial n-1 (Tn-1)*	1,35	3.71	588	.062	.096	0.56	0.62	.458	.016

*Trial n (Tn)*	1,35	0.75	1827	.391	.021	**84.62	3.07	<.001	.707**

*O x Tn-1*	1,35	**5.61	913	.024	.138**	1.03	1.51	.316	.029

*O x Tn*	1,35	1.02	905	.319	.028	**8.72	1.80	.006	.199**

*Tn-1 x Tn*	1,35	2.23	841	.144	.060	0.32	0.95	.576	.009

*O x Tn-1 x Tn*	1,35	0.98	1530	.329	.027	0.65	1.33	.425	.018

___________________________________________________________________________________________________________

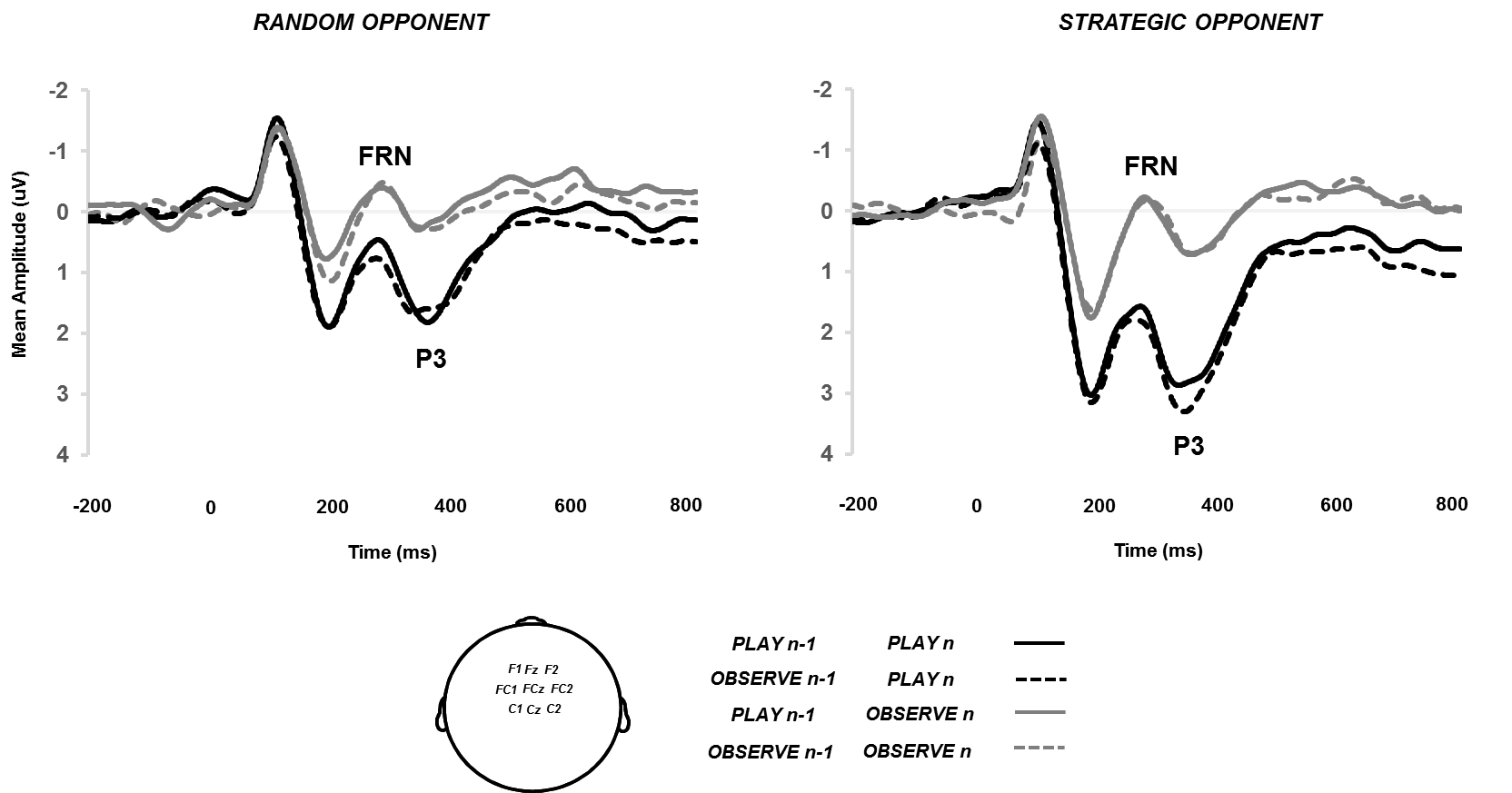

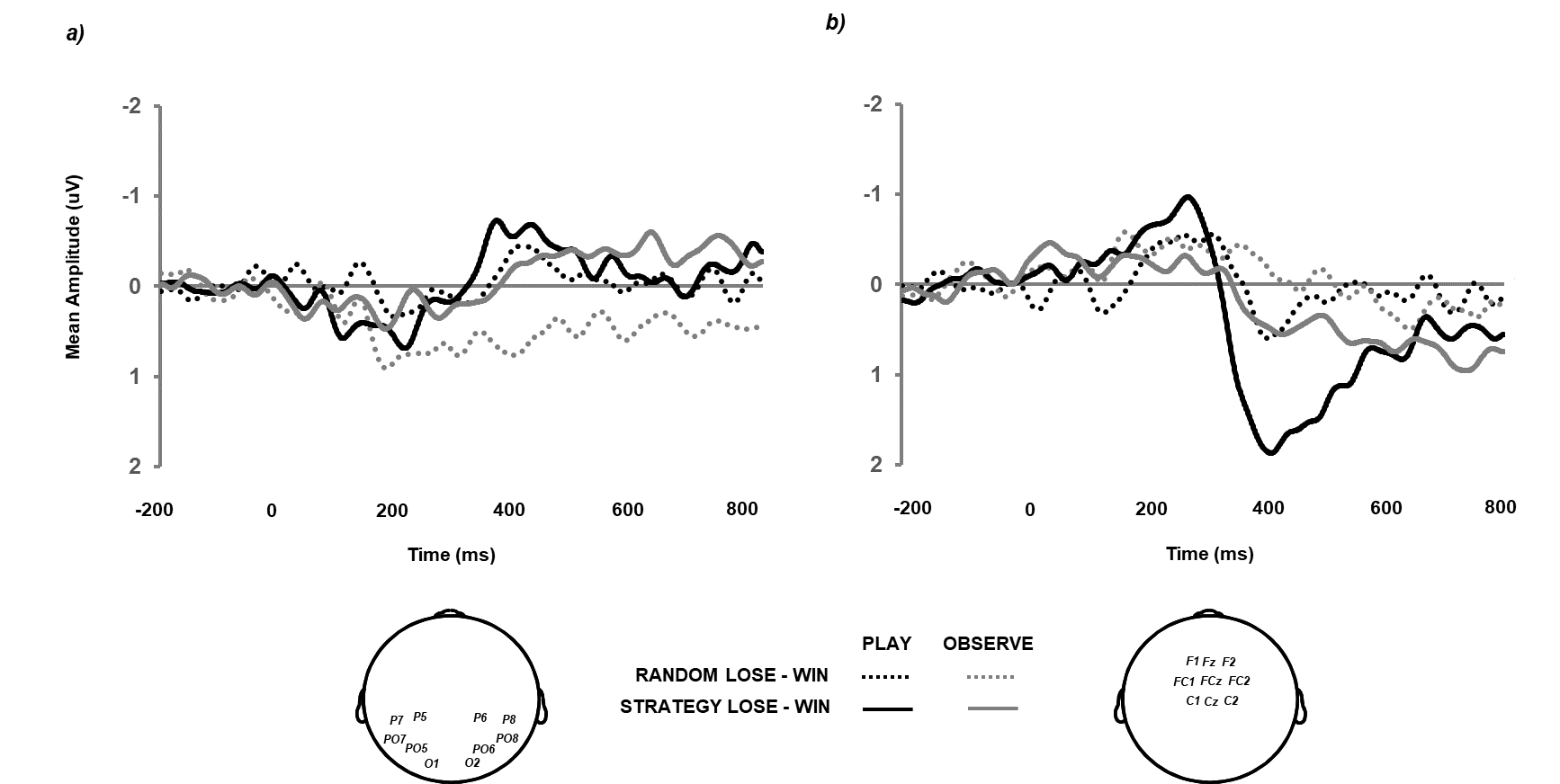

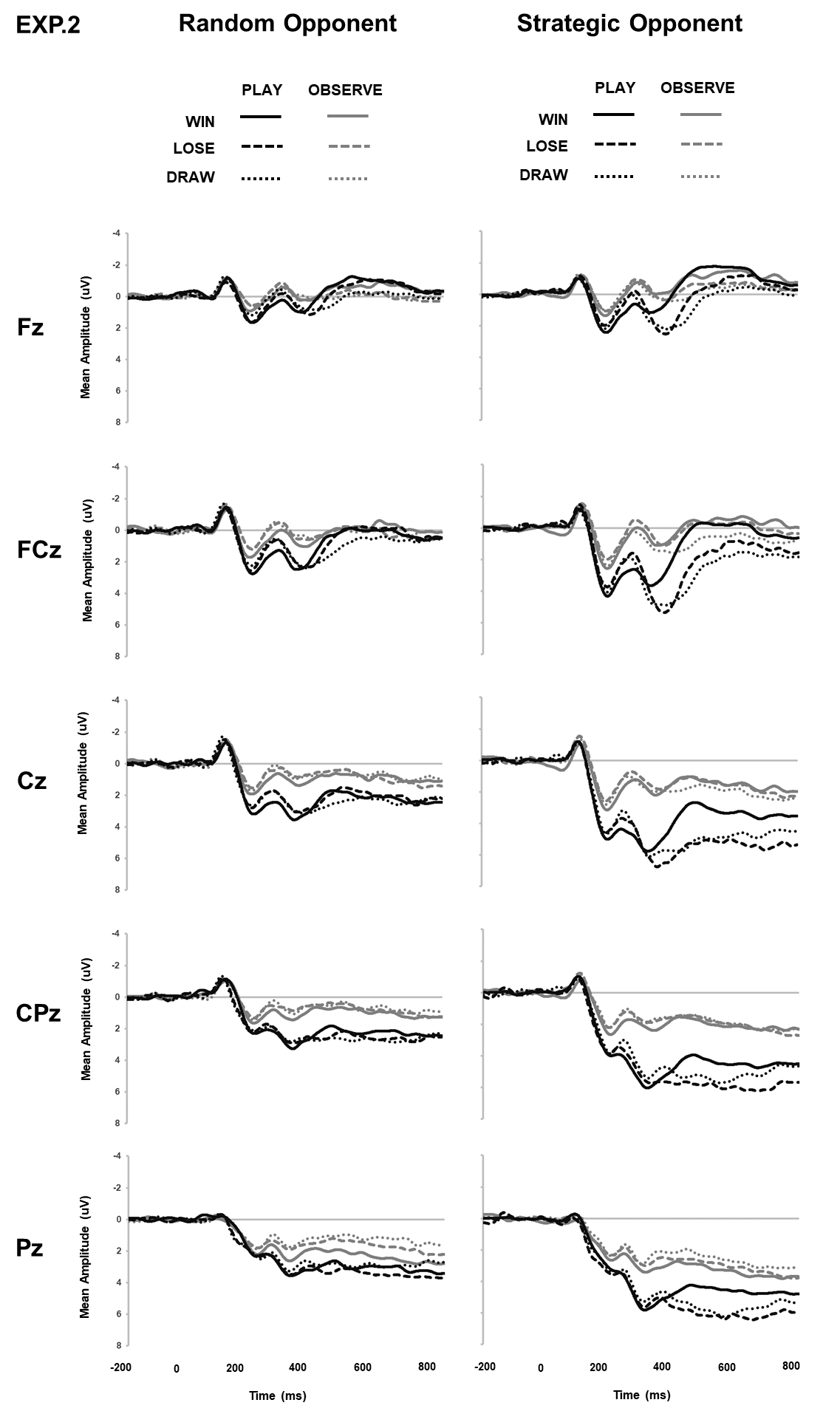

Figure Caption

Supplementary Figure B1. Group-average ERP from 9 fronto-central sites generated by the presentation of trial feedback (win, lose, draw) as a function of the relationship between the type of trial n-1 (play, observe) and trial n (play, observe) for both random and strategic opponents (20 Hz filter applied for presentation).

Supplementary Figure B2. Group-average ERP from a) 10 parieto-occipital and b) 9 fronto-central sites generated by the difference between lose and win trials as a function of trial type (play, observe) and opponent (random, strategic; 20 Hz filter applied for presentation).

Supplementary Figure B3. Group-average ERP generated by the presentation of trial feedback (win, lose, draw) as a function of trial type (play, observe) and opponent (random, strategic) across 5 mid-line electrodes from anterior (Fz) to posterior (Pz) sites (20 Hz filter applied for presentation).
